# The limbic navigation system as a hierarchical RNN

**DOI:** 10.64898/2026.10.19.760224

**Authors:** Zilong Ji, Huiwen Zhang, Krishna Gorantla, Neil Burgess

**Affiliations:** UCL Institute of Cognitive Neuroscience, University College London, London, UK; UCL Queen Square Institute of Neurology, University College London, London, UK

**Author notes:** Equal contribution (Zilong Ji, Huiwen Zhang).

## Abstract

Path-integration-trained recurrent neural networks (RNNs) can reproduce grid-like and head-direction-like representations, but most existing models rely on low-dimensional velocity inputs and do not capture the hierarchical organisation of the brain’s self-localisation circuit. We introduce a hierarchical recurrent neural network (HRNN) that separately processes angular velocity and linear speed to perform simultaneous angular and positional path integration. Trained on these tasks, the HRNN generalises to longer timescales than seen during training and develops neural response properties resembling all the key cell types: angular velocity cells, head direction cells, speed cells, conjunctive and pure grid cells. The learned circuit also recovers key motifs of the navigation system, including attractor dynamics and asymmetric connectivity patterns. Perturbation analyses reveal functional information flow across the hierarchy, consistent with experimental observations. Finally, introducing theta-paced mutual inhibition between two learned angular-velocity subpopulations enables the HRNN to reproduce the bidirectional theta sweeps observed in empirical data. These results establish hierarchical task-driven RNNs as a biologically interpretable framework for linking neural representations, circuit motifs, and population dynamics in spatial navigation, while enabling targeted in silico perturbations and the generation of experimentally testable predictions.

## 1 Introduction

Recurrent neural networks (RNNs) trained with machine learning techniques on cognitive tasks have become a widely adopted tool for neuroscientists to study biological neural circuits [39]. This approach has achieved notable success across multiple domains, including modelling neural activity in visual cortex [20], motor systems [19], and prefrontal cortex [17]. In the domain of spatial navigation, RNNs trained to perform path integration (PI) tasks have been shown to give rise to representations resembling head direction cells [9; 34] and grid cells [8; 1; 28]. Interestingly, beyond reproducing neural activity patterns, these models also exhibit emergent connectivity structures that mirror anatomical motifs observed in biological circuits [9; 7]. This suggests that task-driven RNNs are not merely “black boxes,” but can serve as biologically interpretable models that recapitulate key structural principles of neural systems.

In contrast to mechanistic models—such as continuous attractor neural networks for head direction and grid-cell circuits[27; 41; 5]—which rely on predefined connectivity structures, normative models do not impose such assumptions. Instead, they learn synaptic connectivity directly from task demands, often rediscovering structures consistent with those proposed by mechanistic approaches. As a result, these models provide a complementary perspective, offering a framework to explain why neural circuits take their observed forms across both evolutionary processes and postnatal experience-dependent development.

Despite these advances, current approaches remain limited in their ability to capture the complexity of the spatial navigation circuit. This system depends on the coordinated function of multiple brain regions, each specialized for distinct computational roles. Here we use the term ***limbic navigation system*** to refer broadly to the interconnected circuits that support self-localisation and path integration, including upstream pathways for directional information as well as downstream entorhinal-hippocampal pathways for spatial information (Fig. 1A). For instance, head direction cells in the anterodorsal thalamus and postsubiculum encode orientation information and depend on angular velocity signals from the dorsal tegmental nucleus (DTN) and lateral mammillary nuclei (LMN) [30]. Downstream, grid cells in the entorhinal cortex are thought to continuously track an animal’s position in space [18; 5], relying on upstream head direction and linear speed signals [22; 38]. It is noteworthy that head direction and movement direction are distinct signals, and movement direction, found in a parallel circuit to head-direction [36], is more directly relevant for path integration. Here, we do not dissociate them and instead assume that they are aligned.

**Figure 1:**
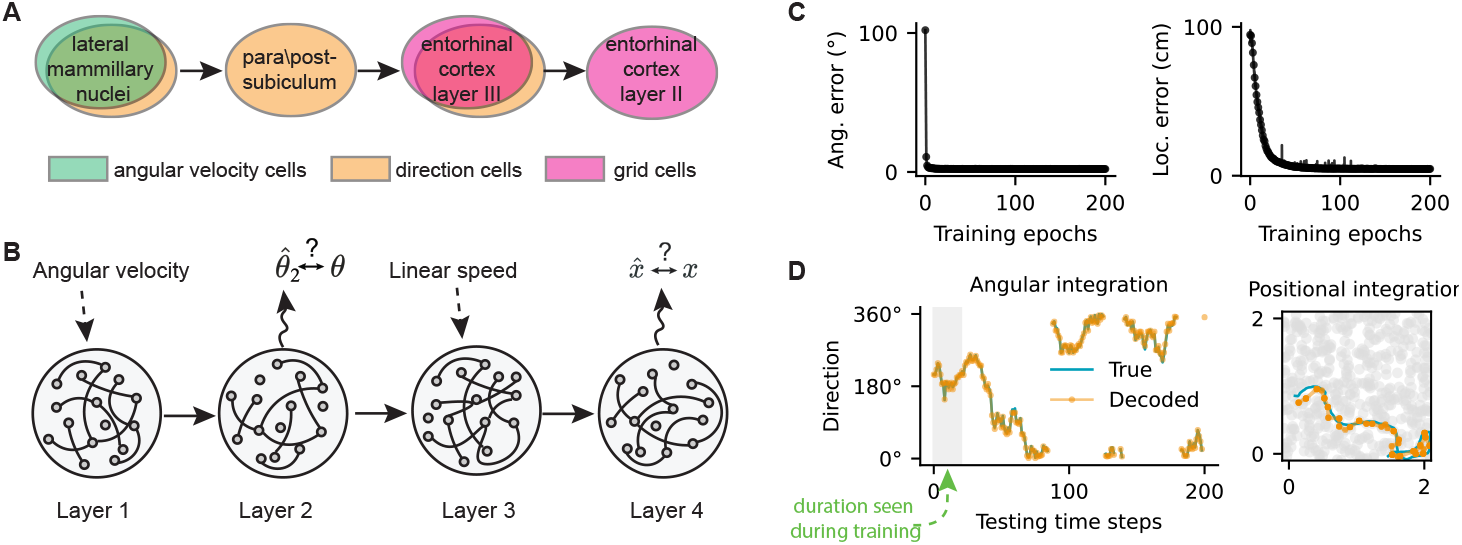
Task-driven hierarchical RNN (HRNN) for simultaneous self-orientation and localisation. (A) Path-integration-related modules in the brain’s limbic navigation system. Different colors indicate different cell types involved in path integration. (B) Schematic of the four-layer HRNN, with each layer corresponding to a module in (A). Angular velocity is provided to the first layer, and the second layer predicts orientation. The third layer receives input from the second layer together with linear speed, and the fourth layer predicts location in 2D space. (C) Test errors across training epochs for angular integration (left) and 2D path integration (right). (D) Path-integration performance after training. Left: angular integration. Right: 2D path integration. The shaded region on the left indicates the time window used during training. The trained HRNN can perform path integration over much longer timescales than those seen during training.

In contrast, most task-driven RNN models rely on a single network trained on a single task with a single optimization objective to capture neural activity patterns, cell types, and circuit motifs, although some studies have begun to explore multi-module or multi-network architectures [34]. Some of these single-RNN models may develop functional submodules during training [40], these emergent structures are often difficult to map onto anatomically distinct areas (but see Cueva et al. [9]). Therefore, they are often limited in their ability to account for the flow of information across interconnected brain regions. Consequently, they fall short of explaining the diversity, organization, and interactions of cell types and circuit motifs distributed across multiple brain regions.

Here, we propose a hierarchical recurrent neural network (HRNN) model trained to perform simultaneous self-localisation of orientation and location. We demonstrate that after training, it recapitulates key organizational principles of the brain’s self-localisation system along three dimensions. First, it reproduces a range of cell types observed across the limbic navigation system, including angular velocity cells [25], head direction cells [31], linear speed cells [15; 13], grid cells [11] and conjunctive direction by grid cells [22]. Second, it captures anatomical motifs, including attractor dynamics reported in the head direction circuit [6], asymmetric synaptic connections [33], aligned directional tuning between functionally connected direction cells and conjunctive grid cells [35], as well as transformations across layers of the medial entorhinal cortex [22]. Third, it captures hierarchical population-level dynamics [29; 16; 35], including recently discovered alternating left-right theta sweeps [35].

The emergence of distinct functional cell types and anatomical motifs enables targeted perturbation experiments in the HRNN, allowing us to investigate the contributions of specific cell populations to self-localisation, in parallel with causal manipulations performed in vivo [38]. Taken together, these results suggest that the HRNN has the potential to serve as a digital twin of the brain’s spatial navigation system, providing a powerful platform for mechanistic investigation and targeted perturbation studies that are difficult to perform in vivo.

## 2 Task-driven hierarchical RNN (HRNN) for self-orientation and localisation

The HRNN consists of four stacked RNNs (Fig. 1B; see Appendix A and Table 1 for detailed parameter settings). Each layer is a “vanilla” RNN [28; 1]. For layer *l*∈{1, 2, 3, 4}, the recurrent update is

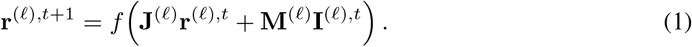

**Table 1:**
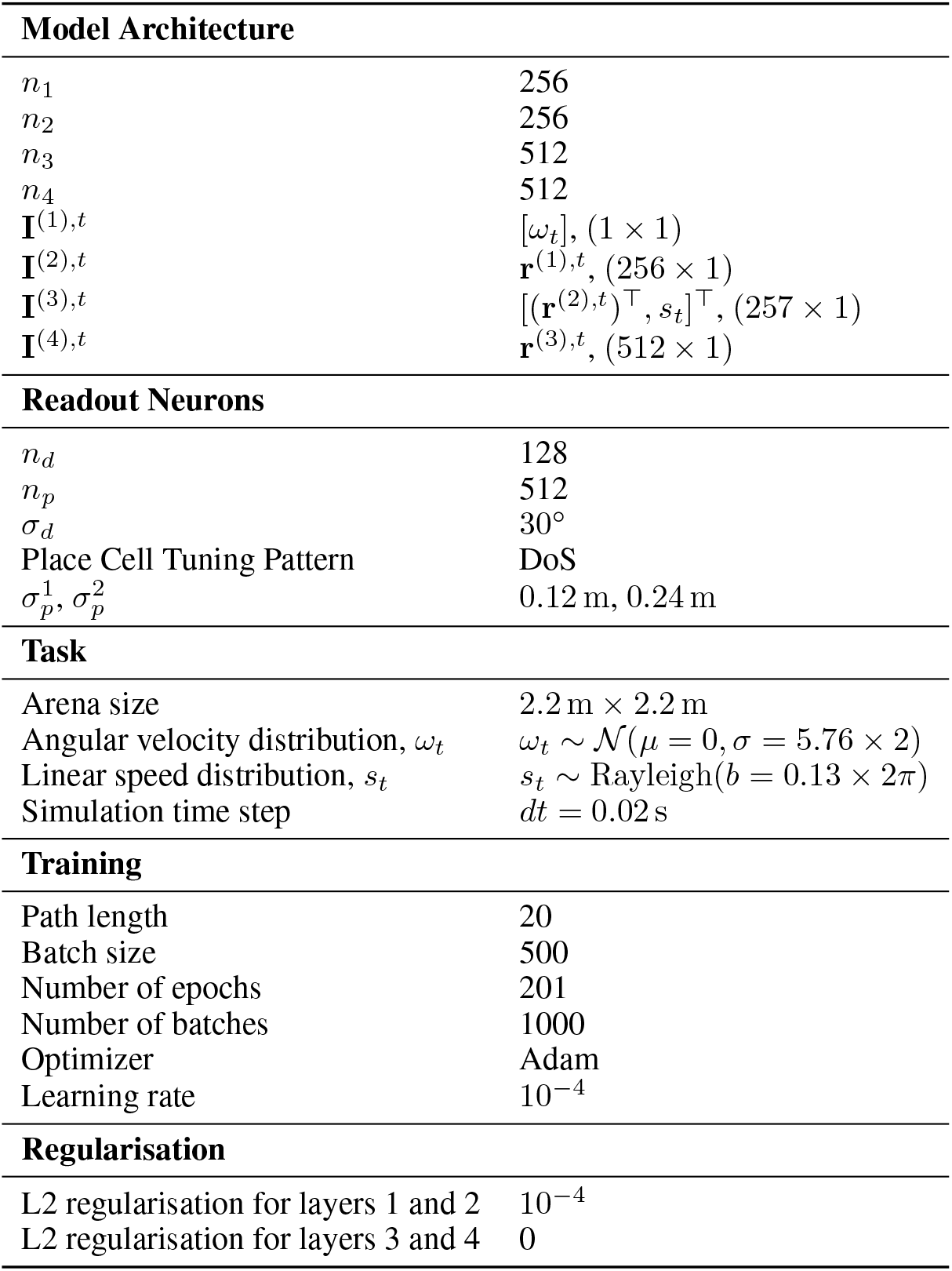
HRNN parameter settings.

| Model Architecture |  |
| --- | --- |
| $n_1$ | 256 |
| $n_2$ | 256 |
| $n_3$ | 512 |
| $n_4$ | 512 |
| $\mathbf{I}^{(1),t}$ | $[\omega_t], (1 \times 1)$ |
| $\mathbf{I}^{(2),t}$ | $\mathbf{r}^{(1),t}, (256 \times 1)$ |
| $\mathbf{I}^{(3),t}$ | $[(\mathbf{r}^{(2),t})^\top, s_t]^\top, (257 \times 1)$ |
| $\mathbf{I}^{(4),t}$ | $\mathbf{r}^{(3),t}, (512 \times 1)$ |
| Readout Neurons |  |
| $n_d$ | 128 |
| $n_p$ | 512 |
| $\sigma_d$ | $30^\circ$ |
| Place Cell Tuning Pattern | DoS |
| $\sigma_p^1, \sigma_p^2$ | 0.12 m, 0.24 m |
| Task |  |
| Arena size | $2.2 \text{ m} \times 2.2 \text{ m}$ |
| Angular velocity distribution, $\omega_t$ | $\omega_t \sim \mathcal{N}(\mu = 0, \sigma = 5.76 \times 2)$ |
| Linear speed distribution, $s_t$ | $s_t \sim \text{Rayleigh}(b = 0.13 \times 2\pi)$ |
| Simulation time step | $dt = 0.02 \text{ s}$ |
| Training |  |
| Path length | 20 |
| Batch size | 500 |
| Number of epochs | 201 |
| Number of batches | 1000 |
| Optimizer | Adam |
| Learning rate | $10^{-4}$ |
| Regularisation |  |
| L2 regularisation for layers 1 and 2 | $10^{-4}$ |
| L2 regularisation for layers 3 and 4 | 0 |

Here, 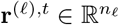 is the activity vector of layer *l* at time *t*, and 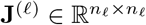 is its recurrent weight matrix. The feedforward input is 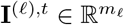, and its weight matrix is 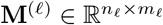. Specifically, **I**^(1),*t*^ = [*ω*_*t*_], **I**^(2),*t*^ = **r**^(1),*t*^, **I**^(3),*t*^ = [(**r**^(2),*t*^)^⊤^, *s*_*t*_]^⊤^, and **I**^(4),*t*^ = **r**^(3),*t*^, where *ω*_*t*_ and *s*_*t*_ are angular velocity and linear speed, respectively. Thus, (*m*_1_, *m*_2_, *m*_3_, *m*_4_) = (1, *n*_1_, *n*_2_ + 1, *n*_3_). The function *f* (*·*) is a ReLU nonlinearity, ensuring that neural activity remains non-negative.

RNN layer 2 outputs the activities of a group of cells coding for orientation, i.e., direction cells:

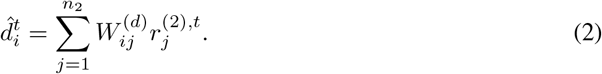

Here, 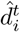is the predicted activity of direction cell *i, n*_*d*_ is the number of direction cells, and 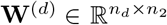 is the direction-readout matrix. Layer 4 outputs the activities of a group of cells coding for location, i.e., place cells:

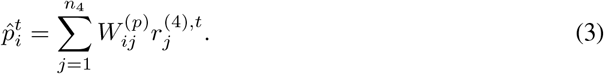

Here, 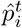 is the predicted activity of place cell *i, n*_*p*_ is the number of place cells, and 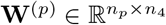 is the place-readout matrix.

The network is trained to simultaneously reconstruct direction cell activities from layer 2 and place cell activities from layer 4:

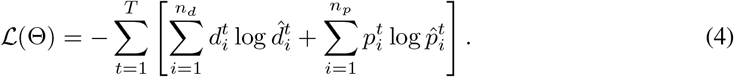

Here, 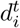 and 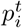 denote the target direction- and place-cell activities encoding the ground-truth direction and position, while 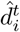 and 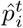 are the corresponding predictions. The two terms are the cross-entropy losses for the layer-2 direction readout and the layer-4 place readout, respectively. The parameter set Θ contains all recurrent, feedforward, and readout weights (see Table 1). We minimise *L*(Θ) using gradient descent while continuously predicting orientation and position over *T* = 20 time steps; prediction errors decrease rapidly during training (Fig. 1C).

After training, the network successfully predicts direction and position trajectories that closely overlap with the ground truth (Fig. 1D). Notably, although it is trained only on short-duration sequences, the network can perform angular and positional path integration over much longer timescales.

## 3 HRNN captures multiple neural activity patterns

After training converged, we examined the spatial tuning of units in the HRNN and found a variety of cell types with distinct tuning properties that closely resemble those reported in the experimental literature (Fig. 2) [31; 25; 29; 24; 11; 22; 15].

**Figure 2:**
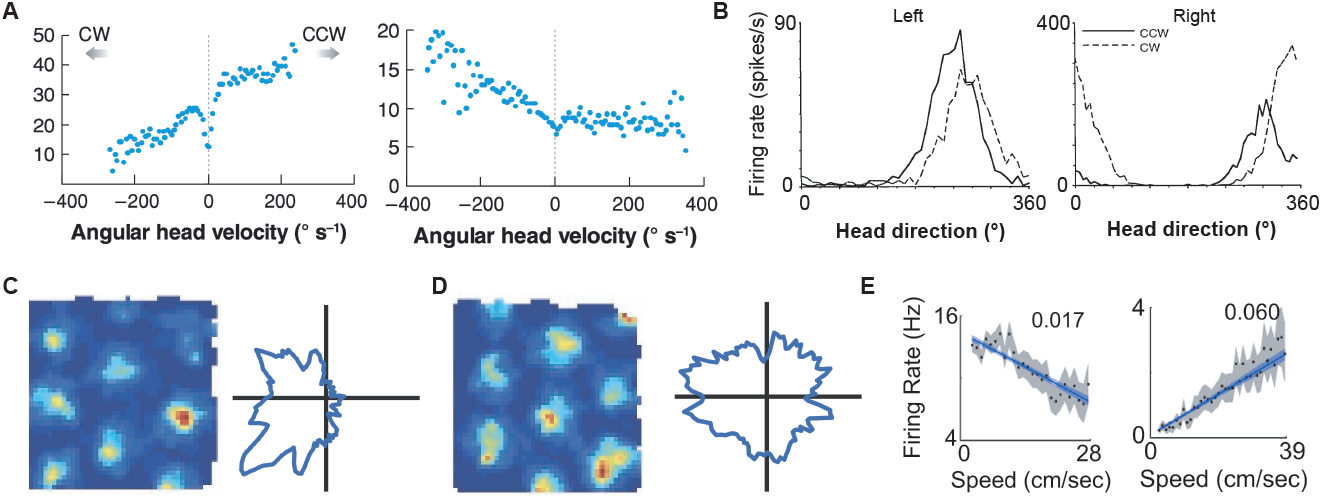
Experimentally identified cell types in the limbic navigation system for self-orientation and localisation. (A) Two angular velocity (AV) cells, adapted from [2]. (B) Two head direction cells showing different firing rates as a function of turning direction, adapted from [29]. (C) A conjunctive grid cell, adapted from [4]. (D) A pure grid cell, also adapted from [4]. (E) Two linear speed cells, one negatively tuned (left) and one positively tuned (right) to linear speed, adapted from [13].

### Angular velocity cells

We first showed the emergence of angular velocity (AV) cells in the first two layers (Fig. 3A). Cells were classified as AV-tuned if they satisfied two criteria: an absolute AV tuning index greater than 0.2 and a linear-fit *R*^2^ greater than 0.5 (see Appendix D). Using these criteria, most of the cells in layer 1 showed AV tuning (218/256), whereas a relatively smaller population in layer 2 showed AV tuning (158/256; Fig. 3B; Fisher’s exact test, odds ratio = 3.56, p = 2.3 *×* 10^*−*9^; see Fig. S8 for all AV tuning maps). This result is consistent with the decrease in AV cells along the head-direction circuit, with a higher proportion in the lateral mammillary nuclei (LMN; [29]) than in the downstream postsubiculum (PoS; [25]).

**Figure 3:**
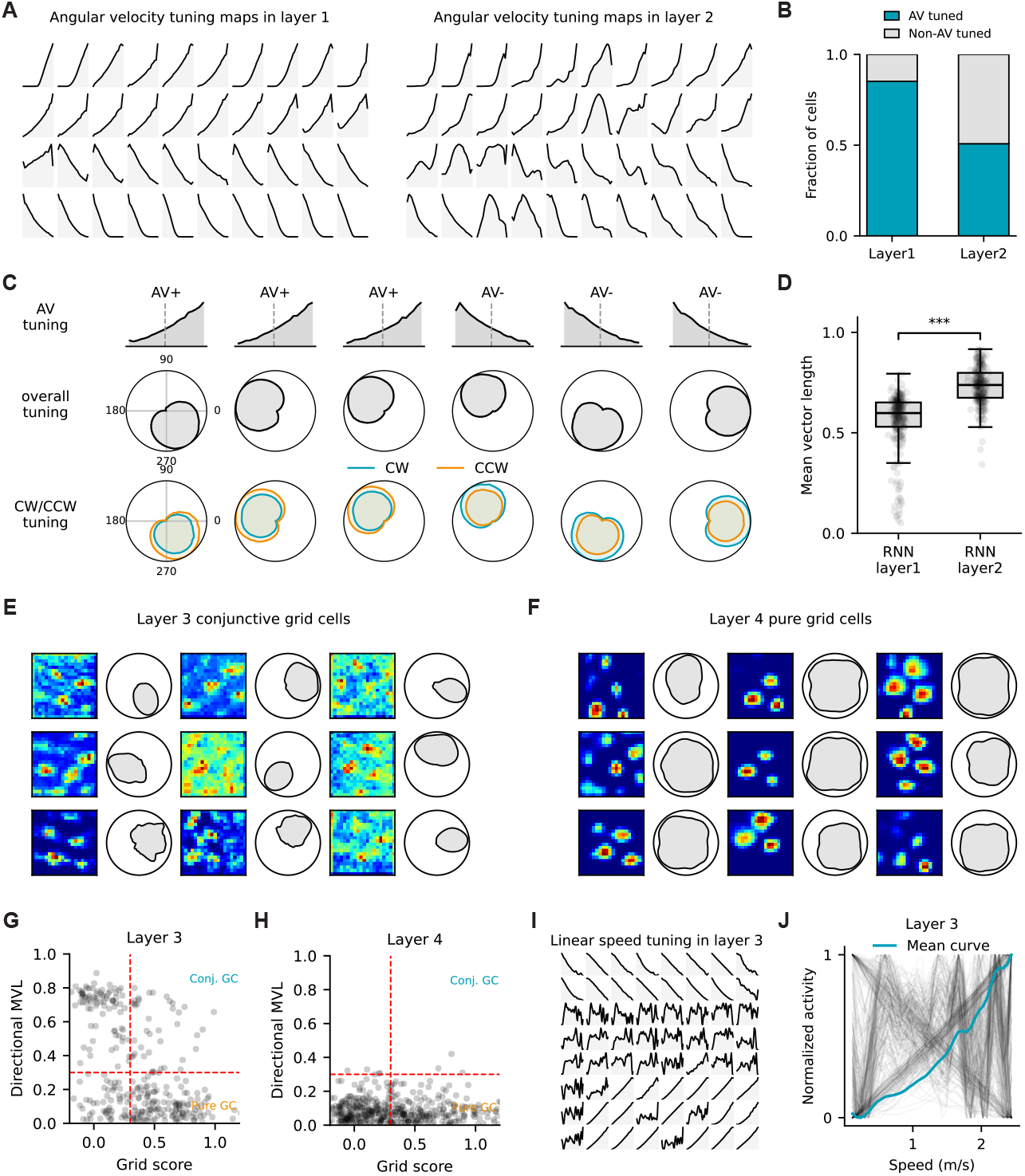
HRNN recapitulates multiple neural activity patterns observed in the self-localisation system. (A) Example angular velocity (AV) tuning cells in layers 1 and 2, respectively. (B) AV tuning strength in layers 1 and 2, showing a significant decrease in AV tuning strength in layer 2 (Fisher’s exact test, odds ratio = 3.56, p = 2.3 *×* 10^*−*9^). (C) Examples of conjunctive AV tuning (row 1) and head-direction tuning (row 2), with clockwise (blue) and counter-clockwise (orange) directional tuning plotted separately. Positive AV-tuned cells preferentially fire in the counter-clockwise (CCW) direction, whereas negative AV-tuned cells preferentially fire in the clockwise (CW) direction. (D) Directional tuning is stronger in layer 2 than in layer 1, reflected by the directional mean vector length, with Mann-Whitney *U* = 6.6*×* 10^3^, *p* = 1.2 *×*10^*−*53^. (E) Conjunctive grid-by-direction tuning cells in layer 3. (F) Pure grid cells in layer 4. (G) Distribution of grid scores and direction mean vector lengths for all active cells in RNN layer 3. (H) Same as (G) but in layer 4. (I) Linear speed cells in layer 3, including both positive and negative speed-tuned cells. (J) Linear speed tuning curves for all active cells in layer 3 overlaid in one plot. The blue line denotes the average linear speed tuning curve.

Furthermore, AV cells can be divided into two subgroups (Fig. 3A&C): one increases the firing rate for counter-clockwise (CCW) turns (angle increases from 0° to 360° with a positive slope; AV+), and the other increases the firing rate for clockwise (CW) turns (angle decreases from 360° to 0° with a negative slope; AV-). The numbers of cells in these two groups are balanced. Knocking out one group causes the directional representation to drift continuously in the opposite direction, yielding an experimentally testable prediction (see Section 5 below).

### Head direction cells

Most of the cells in the first two layers are also conjunctively tuned to directional information (Fig. 3C; see also Fig. S8 for all directional tuning maps). Furthermore, AV+ cells show preferred firing for CCW turns while AV-cells show preferred firing for CW turns. Interestingly, directional tuning is stronger in layer 2 compared to layer 1 (Fig. 3D; median mean vector length (MVL) [IQR]: 0.74 [0.67, 0.80] in layer 2 vs. 0.60 [0.53, 0.65] in layer 1; Mann-Whitney *U* = 6.6 × 10^3^, *p* = 1.2 *×* 10^*−*53^; see Appendix E for calculation). This result recapitulates the stronger directional tuning observed in later hierarchies of the direction circuit than in earlier hierarchies in the rodent brain [3].

### Conjunctive and pure grid cells

In the HRNN, layer 3 receives directional input from layer 2 together with linear speed input and relays this combined signal to layer 4, which is trained for 2D path integration. Cells in layer 3 exhibited tuning maps resembling conjunctive direction-by-grid cells (Fig. 3E; see Appendix F for calculation details), whereas cells in layer 4 resembled pure grid cells (Fig. 3F). This recapitulates empirical findings that mEC layer-3 grid cells show conjunctive directional tuning, whereas mEC layer-2 grid cells downstream are not [11; 22]. The HRNN recapitulates this pattern without explicit constraints (Fig. 3G,H). Using a grid-score threshold of 0.3 and a directional mean-vector-length threshold of 0.3, we identified 33/512 conjunctive grid cells in layer 3 and only 3/512 in layer 4. We also found a large population of cells with weak grid and head-direction tuning (55/512 in layer 3 and 242/512 in layer 4; Fig. S9), which may still be important for accurate position readout because a single grid module alone yields ambiguous location estimates.

### Speed cells

Because linear speed input was provided to layer 3, we also observed speed cells in layers 3 and 4 (Fig. 3I; Fig. S9), consistent with experimental reports [15; 13; 14]. In layer 3, many cells either increased or decreased their firing rates approximately linearly with running speed, although the majority showed positive speed tuning, as reflected in the mean tuning curve (Fig. 3J). Both positive and negative speed tuning have been observed empirically [13]. In layer 4, by contrast, speed tuning was often nonlinear rather than strictly linear, with many cells reaching saturation before intermediate speeds were attained (Fig. S9), again consistent with experimental data [13]. This layer-specific difference in speed tuning yields an experimentally testable prediction.

In summary, the HRNN recapitulates many experimentally identified cell types in the limbic navigation system. Importantly, these cell types are rarely selective for only a single variable such as direction or location. Instead, they are often conjunctively tuned to multiple features. This supports the view that navigation relies on multiplexed neural codes [12].

## 4 HRNN captures anatomical motifs

Here we further showed that HRNN recapitulates anatomical motifs reported in the literature [27; 33; 6; 35].

### 4.1 Attractor dynamics and asymmetric connections for directional coding in layer 1 and 2

Population activity in the head-direction circuit has been shown to lie on a one-dimensional ring-attractor manifold embedded in a high-dimensional neural state space [6]. We found that both layers 1 and 2 emerge ring-attractor dynamics (Fig. 4A). We next examined the flux of network states on and off the manifold using the analysis proposed by Chaudhuri et al. [6] (see Appendix G for more details). This analysis allowed us to test whether states that were temporarily perturbed away from the manifold would return to it, providing evidence for attractor dynamics. We found approximately zero net flux along the manifold, consistent with the isotropic distribution of flow directions along the ring. In contrast, states away from the manifold showed larger net fluxes directed back toward it, consistent with the attractor acting as a low-energy basin (Fig. 4A).

**Figure 4:**
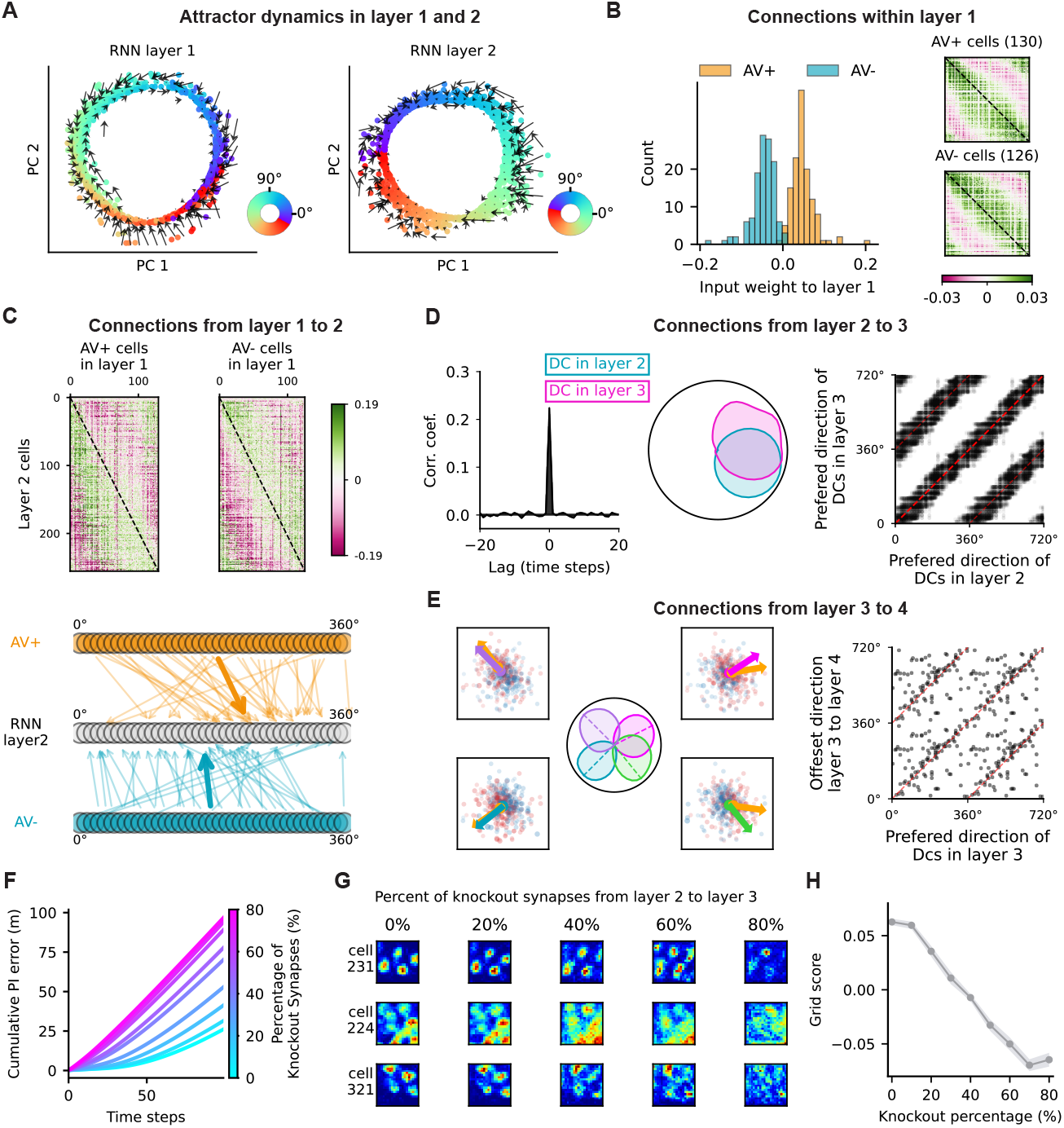
HRNN recapitulates anatomical motifs. (A) Ring-attractor dynamics in layers 1 and 2. Each dot shows the network state projected onto the first two principal components and colored by movement direction. Dark arrows indicate that off-manifold states are driven back toward the attractor manifold. (B) Left: distribution of learned input weights to layer 1, with connections to AV+ and AV-cells shown in orange and blue. Right: asymmetric recurrent connectivity matrices, ordered by preferred direction, for AV+ (top) and AV- (bottom) cells. (C) Top: asymmetric feedforward connectivity from AV+ (left) and AV- (right) cells to layer 2 neurons. Bottom: strongest synaptic connections from AV+/AV-cells to layer 2 cells, highlighting the same asymmetry. (D) Left: cross-correlation of firing sequences for one layer 2 neuron (blue) and one layer 3 neuron (pink); a peak near zero lag indicates co-activation. Middle: directional tuning of the same pair. Right: preferred directions of all functionally connected layer 2-layer 3 pairs, repeated for two cycles for better visualisation. (E) Left: four example layer 3 direction cells. Arrows (not in the orange color) indicate preferred firing direction; red and blue dots mark the spatial firing center of positively and negatively connected layer 4 cells, with dot size proportional to connection strength; the orange dot marks the weighted mean spatial offset. Right: preferred direction of layer 4 cells versus the offset direction of the mean spatial map center in layer 4. (F) Cumulative path-integration error increases as the proportion of pruned synapses from layer 2 to 3 increases. (G) Hexagonal firing fields of three example grid cells in layer 4 progressively degrade with increasing pruning of layer 2-to-layer 3 synapses. (H) Mean grid score across all 4 cells decreases as the pruning percentage increases.

Many previous mechanistic models of angular integration share a common architecture, although they differ in implementation details [27; 26; 41]. This architecture includes (1) cells that encode angular velocity, (2) cells with conjunctive tuning to angular velocity and head direction, and (3) asymmetric synaptic connectivity that shifts the activity bump around the ring. We have already shown the first two features above and here further show that the first two layers also develop asymmetric synaptic connectivity. Specifically, an AV+ cell with a given preferred direction formed its strongest connection with an AV+ cell whose preferred direction was slightly shifted counterclockwise (Fig. 4B). By contrast, an AV-cell connected most strongly to an AV-cell whose preferred direction was slightly shifted clockwise. This learned asymmetry allows angular velocity inputs of opposite signs to shift the activity bump on the ring manifold in opposite directions. Although this pattern is difficult to identify in rodents, it has been elegantly demonstrated in the fly head-direction system [33] and was also captured by a single-layer RNN trained for angular integration [9].

Furthermore, AV+ and AV-cells projected asymmetrically to direction cells in layer 2, but with the opposite shift direction (Fig. 4C). This connectivity pattern has previously been hypothesized in mechanistic models that hard-code this feature to implement angular integration [27; 26; 41]. Here, we show that in a network trained for angular integration, this type of asymmetric connectivity emerges naturally.

### 4.2 Direction-dependent shift for spatial coding in layer 3 and 4

Similar to angular integration, 2D path integration requires the network to continuously update position relative to a starting point using momentary speed and direction. We next examined how this information is conveyed through the learned motifs in the last two layers. We found that direction cells in layer 2 were functionally coupled to direction cells in layer 3 with a similar preferred direction, some of which were conjunctive grid cells as reported in Vollan et al. [35] (Fig. 4D). To identify co-active pairs, we computed the cross-correlation of their firing sequences; pairs with a peak at zero lag were classified as functionally connected. Among 256 direction cells in layer 2 and 105 direction cells in layer 3 (mean vector length, MVL *>* 0.5), we identified 7337 functionally connected pairs whose preferred directions were strongly aligned (Fig. 4D).

Because direction cells in layer 3 are also spatially tuned, we next asked how their activity drives position updates in layer 4. We found that a direction cell in layer 3 tends to activate layer 4 cells ahead of the same direction when the agent moves along that direction, and suppress layer 4 cells on the opposite direction (Fig. 4E). We further computed the center of each layer 4 spatial field and weighted it by the absolute value of the feedforward connection from a probe layer 3 cell (see Appendix H for the detailed calculation). Across 103 identified direction cells in layer 3, the resulting mean spatial offset in layer 4 was aligned with the probe cell’s preferred firing direction (Fig. 4E).

These results suggest that movement in a given direction activates direction cells in layer 2, which in turn recruit layer 3 direction cells with similar preferred directions. These cells then shift population activity in layer 4 along the same direction, thereby continuously updating position from self-motion cues.

### 4.3 Disrupting direction signal impairs grid cell firing

Because the HRNN recapitulates cell types and anatomical connectivity patterns found in the brain, it enables in silico targeted perturbation experiments to test how perturbing one cell type affects the activity and function of others.

Here, we investigated how directional signals influence the generation and function of grid cell activity. Previous experiments have shown that disrupting head direction cells in the anterodorsal thalamic nucleus (ADN), either by lesioning the ADN or by transiently inactivating it with lidocaine, impairs grid cell firing patterns [38]. To test whether the HRNN reproduces this phenomenon, we pruned synaptic connections from layer 2 to layer 3 and evaluated both path-integration performance and grid cell activity. Increasing the degree of pruning led to a more rapid increase in path-integration error over time (Fig. 4F) and progressively degraded grid firing patterns (Fig. 4G,H), consistent with the empirical findings [38].

## 5 Left-right theta sweeps via mutual inhibition between the AV+ and AV-modules in HRNN layer 1

Recent empirical data showed that, during free movement, decoded positions from mEC population activities exhibit rhythmic “theta sweeps,” transiently representing locations extending away from the animal’s current position at theta frequency and alternating between left and right across successive theta cycles (Fig. 5A) [35]. These left-right sweeps in open space are further driven by bidirectional sweeps from side to side of the head axis in the parasubiculum (Fig. 5B).

**Figure 5:**
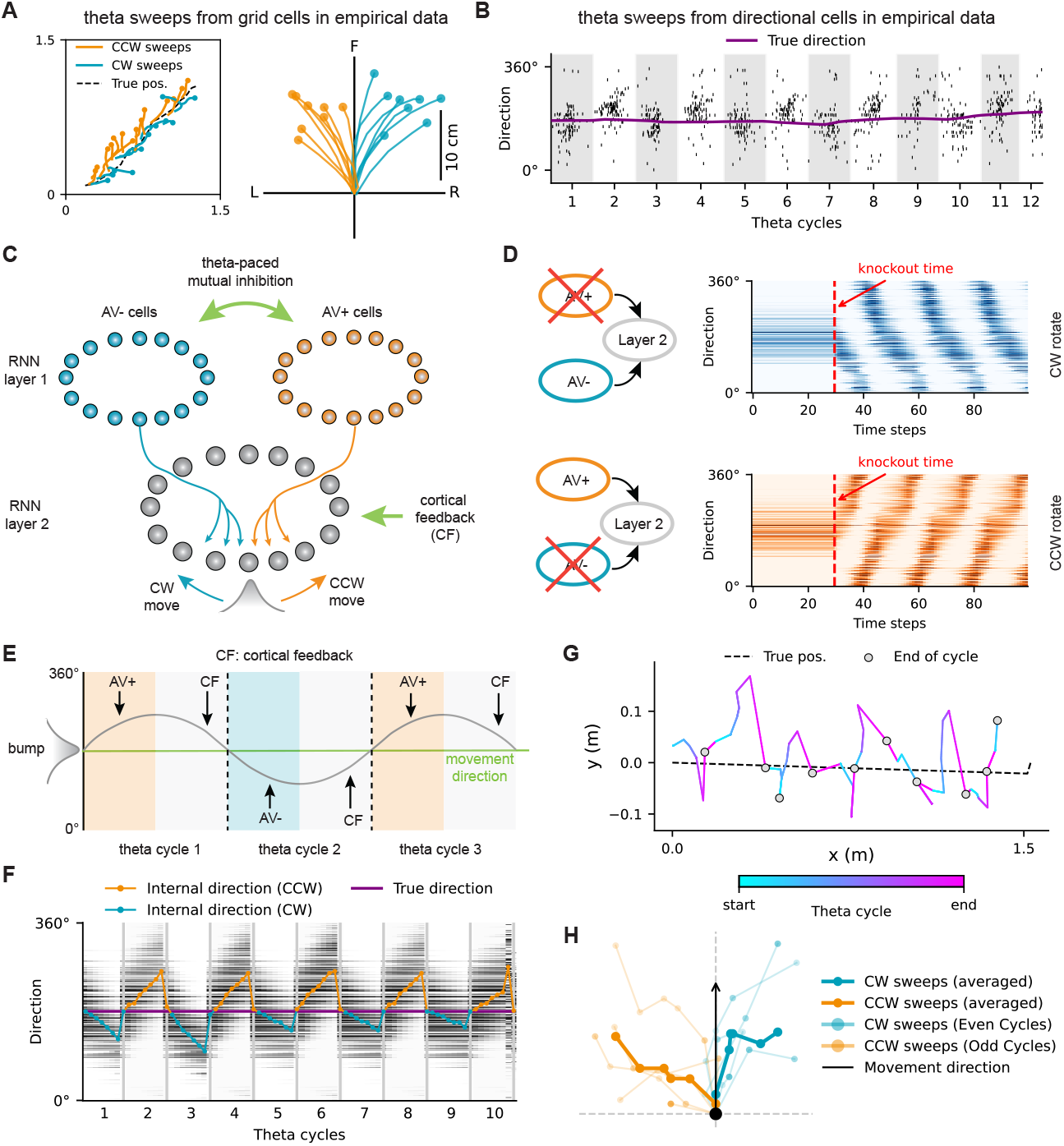
HRNN recapitulates left-right theta sweeps via theta-paced mutual inhibition between the AV+ and AV-cell populations. (A) Left-right theta sweeps decoded from grid cell population activity with data from [35]. Left: left-right sweeps along a straight trajectory. Right: left-right sweeps with initial decoded location recentered at the origin and movement direction pointing to the north. (B) Bidirectional theta sweeps decoded from direction cells in the parasubiculum, with data from [35]. (C) A schematic of the AV+ and AV-populations in layer 1 and the feedforward asymmetric connections from layer 1 to 2. (D) Knocking out AV+ cells (orange) in layer 1 drives the activity bump in layer 2 to drift clockwise, whereas knocking out AV-cells (blue) in layer 1 drives the activity bump in layer 2 to drift counter-clockwise. The red dashed line marks the time of knockout. (E) A schematic of alternatively inhibiting either the AV+ or AV-population across successive theta cycles. At later times in each theta cycle, cortical feedback is introduced to pin the activity bump back to the movement direction. (F) Interleaved inhibition of AV+ and AV-cell populations in layer 1 and the resulting directional sweeps in layer 2 across successive theta cycles. (G) Left-right sweeps in individual theta cycles (from blue to cyan) in layer 4, decoded from the population activity of layer 4 cells. (H) Sweeps in even (blue) and odd (orange) theta cycles after recentering the starting point of each sweep chunk to the origin and reorienting the movement direction to the north, with averaged left and right sweeps plotted as thicker lines.

We have shown that, AV+ cells in layer 1 preferentially project to downstream directional cells in layer 2 whose tuning fields are slightly shifted to the counter-clockwise side of the AV+ tuning fields, whereas AV-cells preferentially project to downstream directional cells whose tuning fields are slightly shifted to the clockwise side of the AV-tuning fields (Fig. 4C and Fig. 5C). Consistent with this connectivity, knocking out AV+ cells causes the activity bump in layer 2 to drift clockwise, whereas knocking out AV-cells causes it to drift counter-clockwise (Fig. 5D).

Based on this mechanism, we alternately inhibited AV+ and AV-cell populations across successive theta cycles and added cortical sensory feedback to pin the activity bump back to the movement direction at the end of each theta cycle (Fig. 5E). This theta-paced mutual inhibition produced bidirectional theta sweeps in layer 2 (Fig. 5F). Moreover, downstream grid cells in layer 4 exhibited alternating left-right theta sweeps (Fig. 5G&H), as observed in empirical data [35].

In summary, these results show that the HRNN trained on path-integration tasks can capture not only anatomical motifs, but also population-level dynamics that arise from these learned motifs. Notably, these dynamics emerge without imposing explicit biophysical constraints on the HRNN.

## 6 Model comparison

To distinguish the contributions of modular input-output organisation and hierarchical information routing, we compared the HRNN with three control architectures (Fig. S2A–D). Here, modularity refers to the assignment of angular-velocity and linear-speed inputs to different modules, with direction and position read out from their respective processing stages. Hierarchy refers to the restriction of inter-module connections to sequential feedforward pathways. The flat RNN receives both inputs and produces both outputs from a single recurrent population. The unstructured HRNN retains four serially connected recurrent layers, but receives both inputs in layer 1 and produces both outputs from layer 4. The fully connected multi-module RNN preserves the HRNN’s input-output arrangement while allowing recurrent connections between all modules. These controls therefore distinguish the effects of specialised input–output organisation from those of constrained inter-module routing.

We conducted both parameter-matched and unit-number-matched comparisons; architectural details, training results and quantitative analyses are provided in Appendix C. Because parameter matching required smaller populations in the fully connected control, we focus here on the unit-number-matched comparison to assess the contributions of modularity and hierarchy independently of differences in population size.

The unit-number-matched comparison showed that the fully connected multi-module RNN reproduced the principal layer-specific trends observed in the HRNN: stronger directional tuning in layer 2 than in layer 1, a higher proportion of conjunctive grid cells in layer 3 than in layer 4, and a higher proportion of pure grid cells in layer 4 than in layer 3 (Table 4). The unstructured HRNN exhibited some of these trends, but did not show a significant layer-wise difference in conjunctive grid-cell fraction. Thus, the layer-specific organisation of neural responses does not require strictly feedforward hierarchy. Instead, these results support modular input–output organisation as an important contributor to the distribution of directional and spatial representations across populations.

We next examined whether the similarities in layer-specific cell properties extended to population dynamics and connectivity motifs. Across ten independent initialisations, the fully connected multi-module RNN had lower layer-1 ring and asymmetric-connectivity scores than the HRNN, despite having identical population sizes and the same input–output arrangement (Table 4). The unstructured HRNN also had lower scores than the HRNN. However, the contribution of hierarchy should not be interpreted as a general requirement for ring-attractor-like dynamics or path integration. The flat RNN also developed ring-like dynamics and asymmetric connectivity, and perturbing inter-layer connections impaired positional path integration in all three multi-layer architectures (Fig. S4E–F).

These comparisons suggest that modular organisation and hierarchical routing make complementary contributions: the former supports layer-specific representations, whereas their combination supports strong ring-like dynamics and organised connectivity asymmetry.

## 7 Discussion

Training RNNs on path-integration tasks has become an established approach for reproducing grid-like firing patterns in the entorhinal cortex [1; 8; 37; 21; 28; 7]; see Dorrell and Whittington [10] for a review. One limitation of this approach, however, is that the emergence of grid-like patterns can depend on specific modeling and training choices, including hyperparameter settings and the form of supervised place-cell targets [23]. Even so, these models remain valuable normative frameworks for understanding why grid-like representations can arise from the computational demands of path integration. Most previous models, however, supplied the network with two-dimensional velocity inputs, typically expressed as allocentric components *v*_*x*_ and *v*_*y*_. While convenient computationally, such a representation is unlikely to be directly available to the brain, because it requires self-motion signals to be transformed into an external coordinate frame, for example relative to the walls of a square environment. In the brain, angular direction and linear speed are represented by distinct cell populations, including head direction cells [31; 36] and linear speed cells [15; 13], without any need for an explicit decomposition of velocity into two orthogonal allocentric components. Therefore, our HRNN separates directional and speed processing, which is more consistent with the biological organization of the navigation circuit.

Angular velocity (AV)-by-direction cells have been reported in the LMN, where they also show hemispheric lateralization [29]. Most direction cells recorded in the left LMN exhibit higher firing rates during counterclockwise (CCW) turns [29], resembling the AV+ cells in layer 1, whereas most direction cells recorded in the right LMN exhibit higher firing rates during clockwise (CW) turns, resembling the AV-cells. This raises the possibility that theta-paced interactions between the two hemispheres could lead to directional sweeps generated in the HRNN. Although the classical head direction circuit is not considered theta-modulated, direction cells in the anteroventral thalamic nucleus and parasubiculum do show theta modulation [32; 16; 35] (the population that encodes movement direction [36] and drives grid cells via conjunctive grid cells [35]). Such theta modulation may support alternating inhibition between lateralized directional populations, providing a possible biological analogue of the artificially imposed theta-paced oscillations between AV+ and AV-cells that generate theta sweeps in the HRNN. Testing this possibility will require future experiments.

Taken together, our results suggest that the HRNN could provide a first step toward digital twins of the brain’s spatial navigation system. By linking task demands, neural representations, circuit motifs, and population dynamics in a single trainable model, the HRNN offers an integrative and predictive framework for understanding navigation circuits as dynamical systems. More broadly, we suggest that the next frontier is not simply the accumulation of more data, but the construction of mechanistic models with minimal hand-coded pre-specification that can generate testable predictions and guide targeted perturbations before costly in vivo experiments are performed.

## Author contributions

Z.J. and H.Z. contributed equally to this work. K.G. contributed to model simulation. All authors discussed the results, and reviewed the manuscript. Z.J. and N.B. jointly supervised the work.

## Code availability

The source code used to generate the results presented in this paper is publicly available at https://github.com/ZilongJi/HRNN4PI.git.

## A Parameter settings

The HRNN was trained with simulated trajectories generated in a two-dimensional arena. All experiments were performed on a single RTX 3080 GPU. At each time step, angular velocity and linear speed were sampled from predefined distributions (Table 1). Each HRNN layer has *n*_*l*_ hidden units and receives **I**^(*l*),*t*^ from the previous layer and/or from external inputs (angular velocity and linear speed). L2 regularisation was applied to the recurrent weights of hidden units in layer 1 and 2. The model architecture, trajectory parameters, readout-neuron parameters, training hyperparameters, and regularisation terms are summarized in Table 1.

The direction readout cells were defined by randomly tiling *n*_*d*_ direction-cell centres across circular angular space from *−π* to *π*. For the head direction *θ*_*t*_ at time *t*, the circular distance between *θ*_*t*_ and the preferred direction of cell *i, u*_*i*_, was computed as

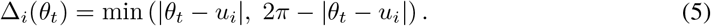

Direction-cell activity was generated using softmax-normalised Gaussian tuning curves,

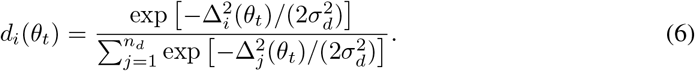

Place cells (readout) were defined by randomly tiling *n*_*p*_ place-cell centres across the 2.2 *×* 2.2 meters two-dimensional environment. For a position **x**_*t*_ at time *t*, the squared distance between **x**_*t*_ and the centre of place cell *i*, **u**_*i*_, was computed as

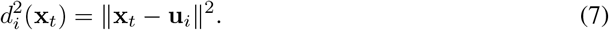

Place-cell activity was generated using a difference-of-Softmax tuning pattern [28]. First, two softmax-normalised Gaussian population activities were computed as

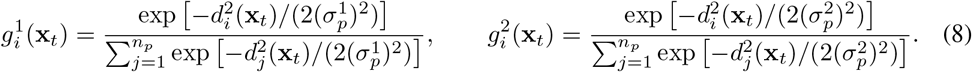

Then, the final place-cell activity for encoding position *x*_*t*_ was obtained from the difference

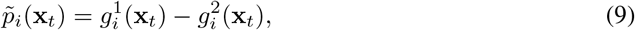

followed by shifting the population activity to be non-negative and normalising it to sum to one across place cells.

## B Parameter sweep for HRNN

To assess whether the reported layer-specific representations and circuit motifs depend on a particular choice of model hyperparameters, we trained 45 HRNN configurations, with training parameters (including learning rate, batch size) the same as reported in Table. 1. Each configuration used an independently sampled random seed. The sweep jointly covered recurrent population sizes, numbers and receptive-field widths of direction and place cells, environment size, and layer-specific recurrent weight decay (Table 2). Thus, the robustness analysis spans substantial variation in both model capacity, task statistics and training strategies, rather than reflecting a single trained network.

**Table 2:** Hyperparameter robustness sweep for the HRNN. We trained 45 configurations for 100 epochs each, using an independently sampled random seed for each configuration. We evaluated all 16 combinations of recurrent layer sizes, and varied input/output population size, tuning widths, environment size, and each layer-specific recurrent weight decay one at a time from the default configuration.

| Hyperparameter | Values tested |
| --- | --- |
| $N_{LMN}, N_{PaS}$ | {128, 256} |
| $N_{Conj}, N_{mEC}$ | {256, 512} |
| Number of direction cells, $N_d$ | {64, 128, 256} |
| Number of place cells, $N_p$ | {256, 384, 512, 640} |
| Direction-cell RF width | {10°, 20°, 30°, 40°, 50°} |
| Place-cell RF width | {0.09, 0.12, 0.15, 0.18} |
| Environment width and height (m) | {1.5, 1.8, 2.0, 2.2, 2.4, 2.6} |
| Recurrent weight decay, each layer | {0, $10^{-5}$, $10^{-4}$, $10^{-3}$} |

**Table 3:** Comparison of the four parameter-matched recurrent architectures. N/A indicates that a layer-wise comparison is not applicable.

| Measure | Flat RNN | Unstructured HRNN | Multi-module RNN | HRNN |
| --- | --- | --- | --- | --- |
| Parameter count | 2.00M | 2.43M | 1.99M | 2.00M |
| Training convergence speed | Slow | Medium | Medium | Fast |
| Fraction of AV cells | 99.7% | 99.0% | 99.7% | 99.6% |
| Fraction of direction cells | 61.1% | 51.9% | 37.6% | 24.1% |
| Fraction of conjunctive grid cells | 4.5% | 2.5% | 1.0% | 8.9% |
| Fraction of pure grid cells | 9.5% | 4.1% | 3.3% | 40.7% |
| More AV cells in Layer 1 than Layer 2 | N/A | No | Yes | Yes |
| Weaker direction tuning in Layer 1 than Layer 2 | N/A | No | Yes | Yes |
| Significant amount of conjunctive grid cells in Layer 3 | N/A | No | No | Yes |
| Significant amount of pure grid cells in Layer 4 | N/A | Yes | Yes | Yes |
| Ring attractor dynamics | Yes | No | No | Yes |
| Asymmetric connections | Yes | No | Yes | Yes |
| Perturbation affects path integration | N/A | Yes | Yes | Yes |

Despite this variation, all configurations learned the path-integration task (Fig. S1A). The resulting representations also preserved the cell types reported in the main text (Fig. S1B): AHV tuning was prevalent in Layer 1, directional tuning was prevalent in Layer 2, conjunctive grid cells were enriched in Layer 3 relative to Layer 4, and pure grid cells were enriched in Layer 4 relative to Layer 3. Importantly, the functional progression of cell types along the hierarchy was also preserved (Fig. S1C,D): paired comparisons across configurations confirmed a decrease in AHV tuning and an increase in directional tuning from Layer 1 to Layer 2, together with the predicted redistribution of conjunctive and pure grid tuning between Layers 3 and 4. This hierarchy of cell types closely resembles the sequence of functional representations reported in rodent navigation circuits [3; 22].

Finally, the circuit-level signatures were similarly robust. Layer 1 activity formed ring-like manifolds across configurations (Fig. S1E), and ring scores exceeded heading-shuffled controls (Fig. S1F). Moreover, preferred-direction-sorted recurrent connectivity consistently showed direction-dependent asymmetry within AV+ and AV*−* subpopulations (Fig. S1G), with asymmetry indices exceeding shuffled preferred-direction controls (Fig. S1H). Together, these results indicate that the hierarchical progression of cell types and the associated Layer 1 dynamical and connectivity motifs are robust across the tested hyperparameter range.

**Figure S1:**
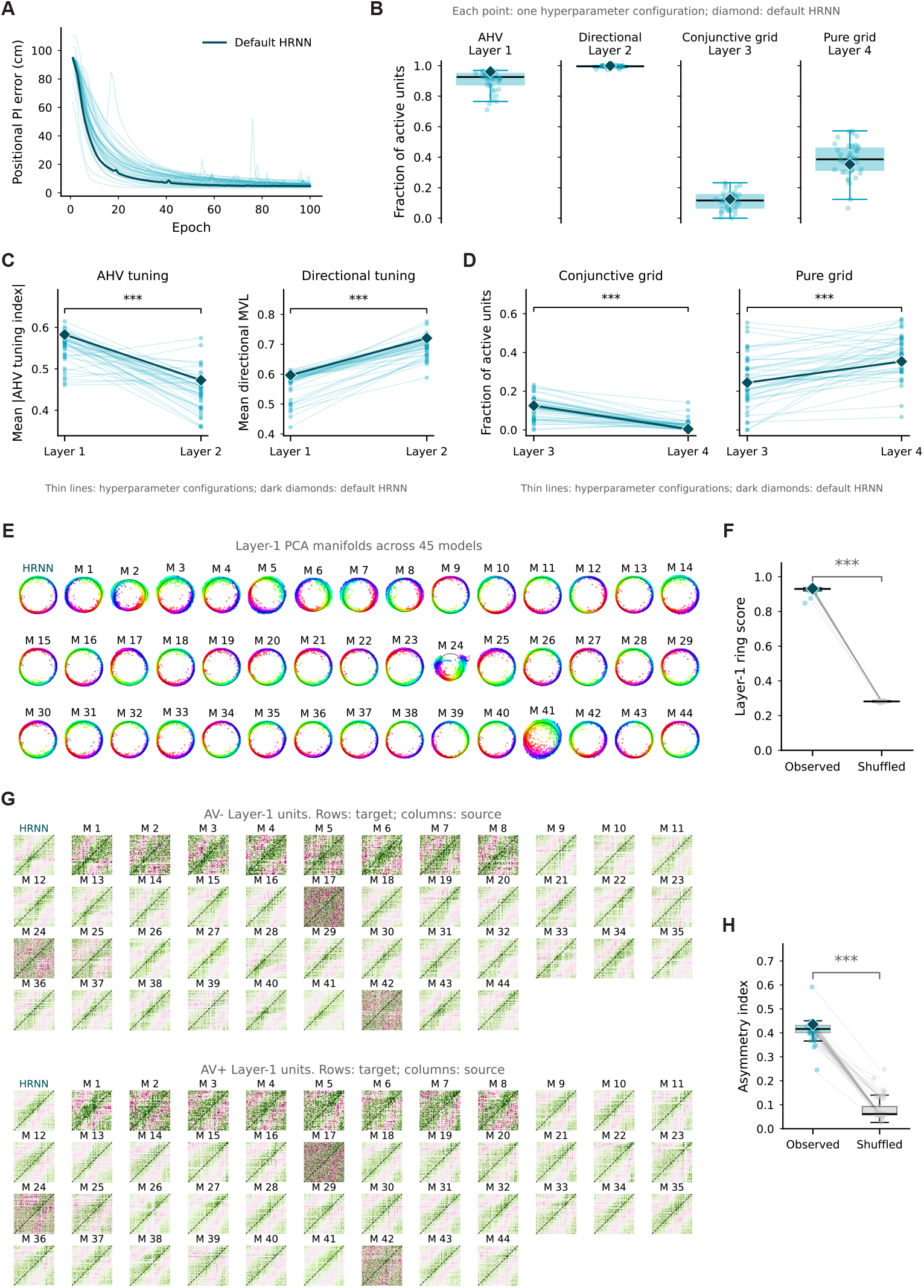
Robustness of HRNN dynamics, cell types, and connectivity across hyperparameter configurations. **A**, Path-integration error during training for 45 HRNN configurations; thin lines denote individual configurations and the dark line denotes the default HRNN. **B**, Fractions of active units exhibiting angular-head-velocity (AHV) tuning in Layer 1, directional tuning in Layer 2, conjunctive grid tuning in Layer 3, and pure grid tuning in Layer 4. **C–D**, Layer-specific tuning transformations across configurations: AHV tuning decreases from Layer 1 to Layer 2, directional tuning increases from Layer 1 to Layer 2, conjunctive grid tuning is enriched in Layer 3 compared to layer 4, and pure grid tuning is enriched in Layer 4 compared to Layer 3. **E**, Two-dimensional PCA projections of Layer 1 activity, reveal ring-like manifolds across configurations. **F**, Layer 1 ring scores exceed heading-shuffled controls. **G**, Recurrent connectivity matrices for AV*−* and AV+ Layer 1 units, show reproducible direction-dependent asymmetric structure. **H**, Preferred-direction asymmetry indices exceed shuffled preferred-direction controls. Diamonds indicate the default HRNN; paired lines link matched observed and shuffled values. \*\*\**p <* 0.001, paired one-sided Wilcoxon test across hyperparameter configurations.

## C Model comparison

The HRNN is biologically constrained. In the rodent brain, angular velocity is processed by the dorsal tegmental nucleus (DTN) and lateral mammillary nuclei (LMN), with directional information read out in the dorsal presubiculum. Linear speed is processed by speed cells in the entorhinal cortex, likely in medial entorhinal cortex (MEC) layer 3, and conveyed to layer 2, where it serves as input to grid cells. Our HRNN was inspired by this hierarchical routing of information.

For computational reasons, we tested whether the proposed HRNN is necessary for efficient path integration and for the emergence of structured neural representations and anatomically interpretable structures.We compared the HRNN with three control architectures trained on identical angular-velocity and linear-speed inputs and the same direction and position targets.

Together, we considered four recurrent architectures with trainable parameters approximately matched (Fig. S2A-D, Table. 3):

1. **Flat RNN**. A single recurrent population receives both angular velocity and linear speed, and jointly predicts direction and position.
2. **Unstructured HRNN**. Four recurrent layers are connected serially, but both angular-velocity and linear-speed inputs are fed into the first layer, and both direction and location are read out from the fourth layer.
3. **Multi-module RNN**. Four recurrent populations receive modality-specific inputs and outputs, but all populations are recurrently connected to one another.
4. **HRNN**. The proposed biologically constrained hierarchy: angular-velocity information is routed through the first two modules, linear-speed and place-related information enter downstream, and outputs are read out at the corresponding stages.

### C.1 HRNN accelerates path-integration learning compared to control architectures

We first compared training dynamics across ten random initializations per architecture. Although all models ultimately reached low positional path-integration error, the HRNN converged substantially faster and with lower between-seed variability than the flat, serial-unstructured, and fully connected multi-module controls (Fig. S2E).

We quantified learning efficiency as the first epoch at which positional error fell below 10 cm. The HRNN reached this criterion consistently within the first few epochs, whereas the other architectures required substantially more training epochs (Fig. S2F). These results show that the advantage is not explained by depth alone or by merely partitioning units into multiple populations; the constrained routing of the HRNN improves optimization efficiency. One potential explanation is that the HRNN first learns directional information in its first two layers (see Fig. 1C). Accurate directional information is then available to layers 3 and 4, allowing them to integrate speed information to estimate position.

**Figure S2:**
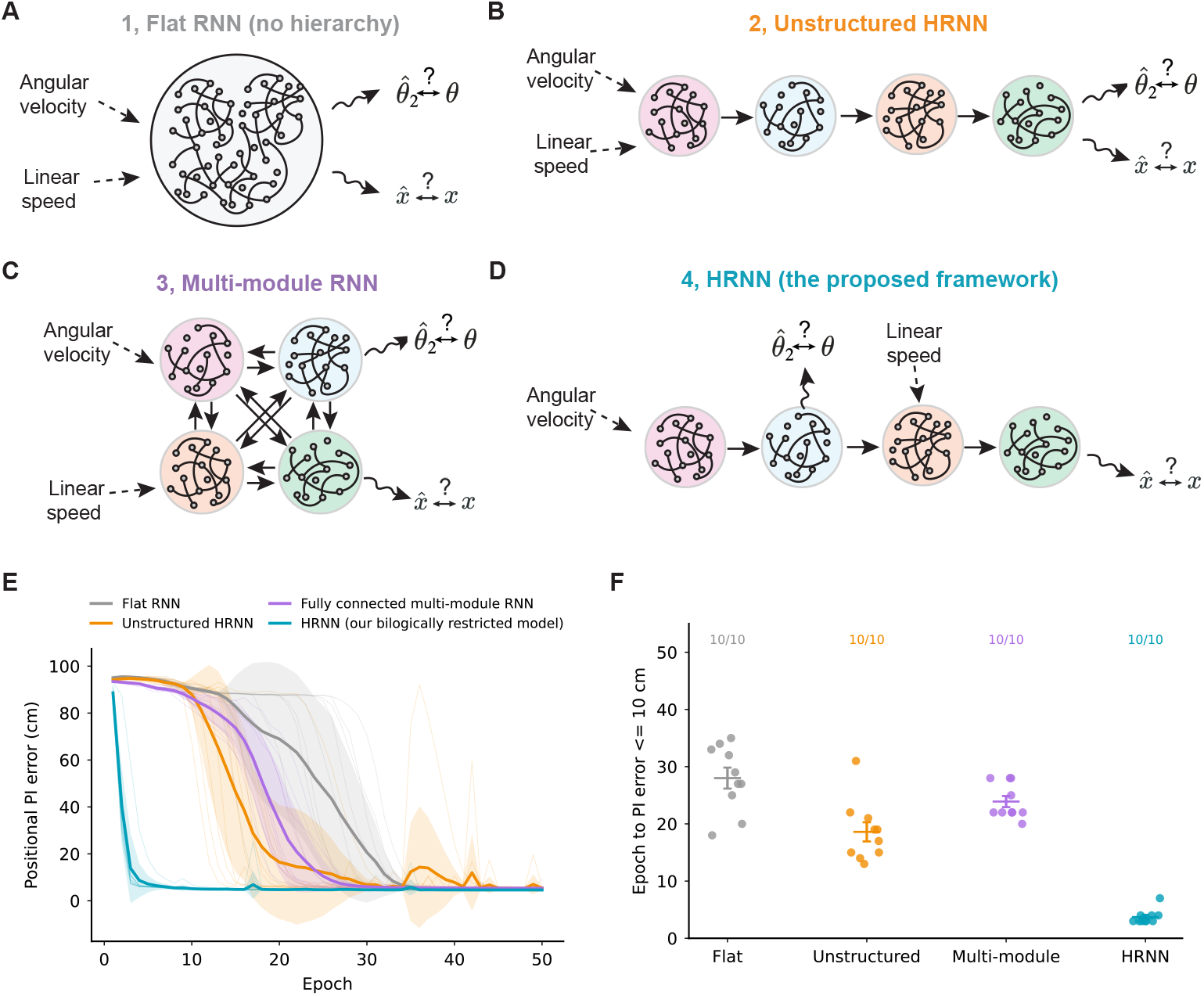
Model comparison reveals that the HRNN accelerates path-integration learning. (A) Flat RNN. (B) Unstructured HRNN. (C) Fully connected multi-module RNN. (D) HRNN. (E) Positional path-integration error across training epochs for ten random initializations of each architecture; faint traces show individual initializations, thick traces show the mean, and shaded regions show variability across initializations. (F) Epoch at which positional path-integration error first reaches 10 cm for each initialization; labels indicate the number of runs (out of ten) reaching this criterion.

### C.2 HRNN promotes layer-specific cell types

**Figure S3:**
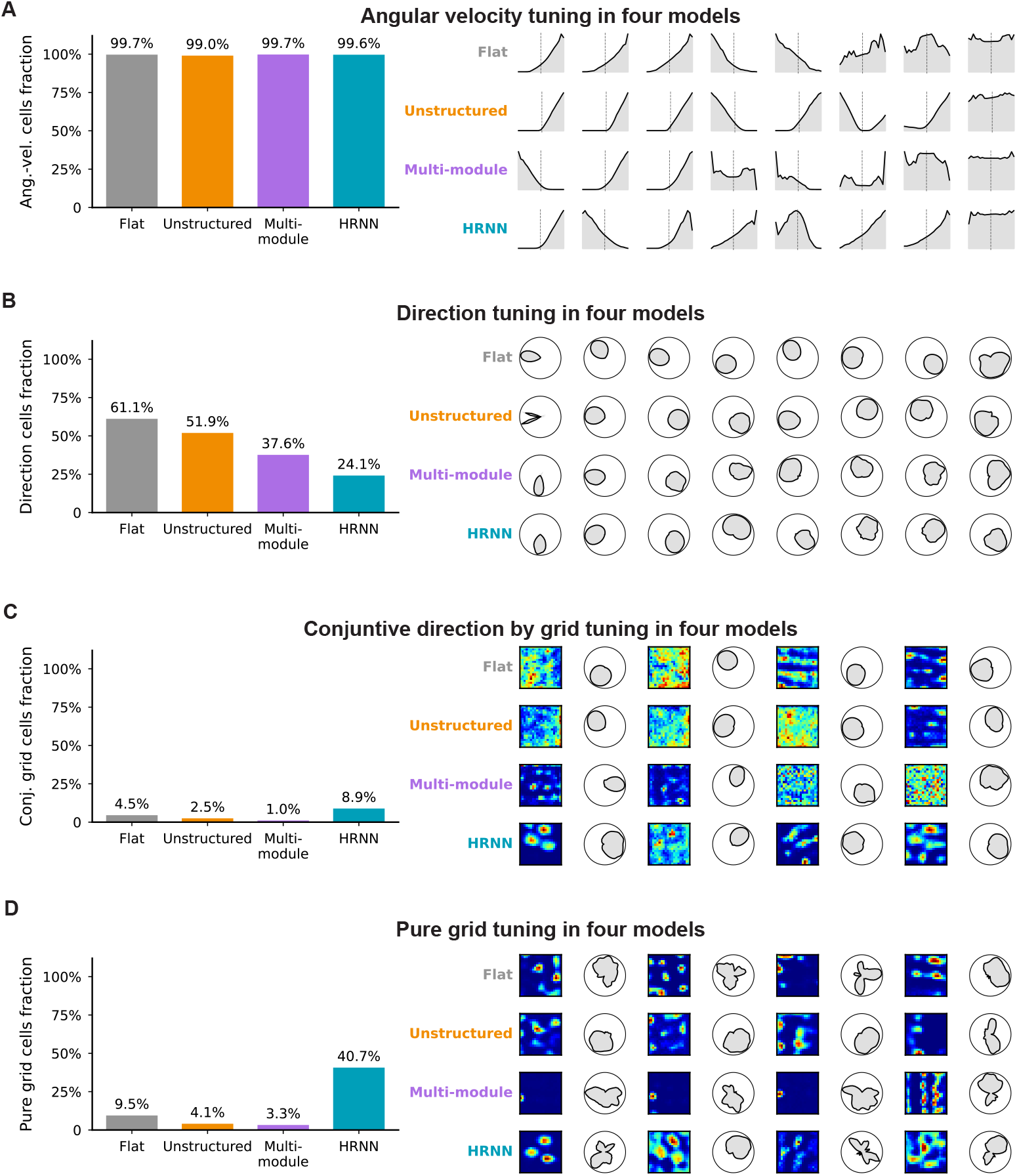
All four recurrent architectures generates biological realistc cell types after training. (A) Fractions of angular-velocity-tuned cells and representative angular-velocity tuning curves for the flat RNN, unstructured HRNN, fully connected multi-module RNN, and HRNN. (B) Fractions of direction-tuned cells and representative directional tuning maps. (C) Fractions of conjunctive direction-by-grid cells and representative spatial firing-rate maps paired with directional tuning maps. (D) Fractions of pure grid cells and representative spatial firing-rate maps paired with directional tuning maps. Bars report the cell fraction in each architecture; colors identify architectures as in Fig. S2.

We next examined whether each architecture developed the cell types expected from its computational organization (Fig. S3). All architectures contained angular-velocity-modulated and directionally tuned units, consistent with the shared path-integration objective. However, their distributions across modules differed markedly. Importantly, only HRNN showed reduced angular velocity tuning and increased diurectional tuning along the hierarchy (Fig. S4A&B), as reported in the biological brain [3].

In the HRNN, early modules were strongly dominated by angular-velocity and directional tuning, whereas downstream modules exhibited conjunctive grid-like and pure grid-like spatial representations. In particular, conjunctive grid cells were enriched in the intermediate spatial module, while pure grid cells were substantially more prevalent in the final module (Fig. S4C&D), as reflected in the bilogical brain [22]. This layer-specific progression was weaker or absent in the flat, unstructured, and fully connected multi-module controls.

These results indicate that task training alone can generate individual tuning motifs, whereas the HRNN promotes their reliable segregation into distinct functional stages, mirroring the organization observed in the biological brain.

### C.3 Only the Flat RNN and the HRNN develop ring attractors in the population activity

**Figure S4:**
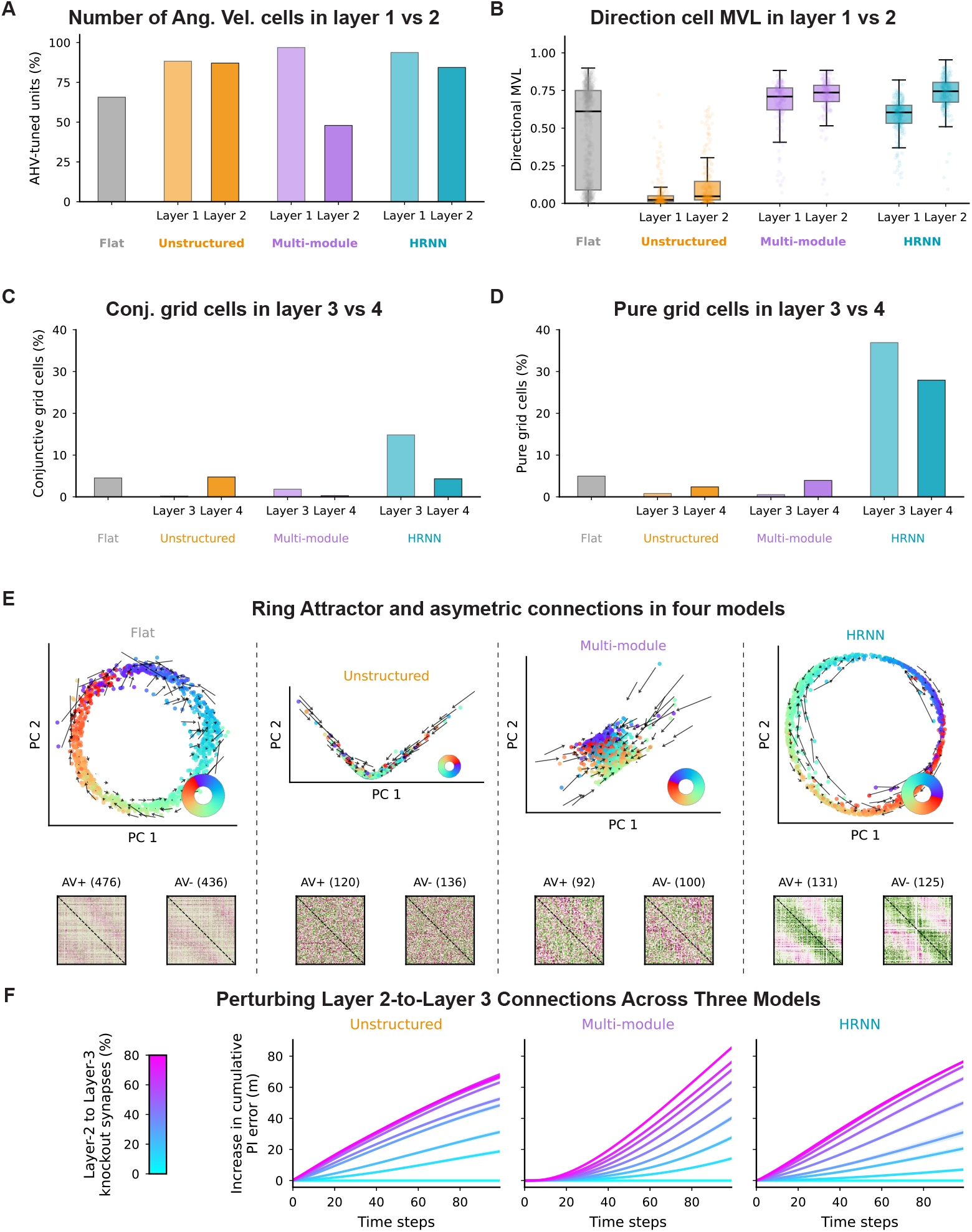
Among the four parameter-matched recurrent architectures, only the HRNN reliably segregates cell types into distinct functional stages, mirroring the organization observed in the biological brain. (A) Proportions of angular-velocity-tuned cells across architectures. (B) Directional tuning strength (mean vector length; MVL) in layers 1 and 2. (C–D) Proportions of conjunctive and pure grid cells in layers 3 and 4. (E) Low-dimensional population dynamics and asymmetric recurrent connectivity between angular-velocity-positive and -negative units. (F) Random knockout of Layer 2-to-Layer 3 synapses increases cumulative positional path-integration error in the unstructured, multi-module, and hierarchical RNNs. Colors indicate the fraction of disrupted synapses.

We then analyzed the geometry and recurrent dynamics of the first recurrent module (Fig. S4). The Flat RNN and HRNN exhibited a circular low-dimensional manifold whose neural trajectories evolved coherently with head direction, consistent with a ring-attractor-like representation. By contrast, the serial-unstructured HRNN and fully connected multi-module RNN showed less organized low-dimensional geometry and less coherent directional flow. The Flat RNN, fully connected multi-module RNN and HRNN showed asymmetric connections, mimicking shifter cells in the brain ((Fig. S4E). Finally, randomly silencing Layer 2-to-Layer 3 synapses increased cumulative positional path-integration error in all three multi-layer models (Fig. S4F).

Together with the layer-specific tuning results, these findings suggest that the HRNN’s constrained routing does more than accelerate learning: it biases the network toward anatomically interpretable connectivity motifs and a sequential transformation from angular-velocity signals to directional, conjunctive, and spatial representations.

### C.4 Unit-matched model comparison

In addition to the parameter-matched model comparison, we also conducted a unit-matched model comparison. The HRNN was compared with the unit-matched unstructured HRNN and the multi-module RNN. All three RNNs contained identical numbers in the four layers (256, 256, 512, and 512). The flat-RNN was excluded for the model comparison because a layer-specific comparison is not applicable to a single-layer architecture.

In summary, the HRNN, with brain-inspired inductive biases, reliably reproduces layer-specific distributions along the navigation hierarchy as well as the relevant ring-like dynamics and asymmetric connectivity motifs. The unit-number-matched multi-module RNN also reproduces aspects of the layer-specific cell-type redistribution, but across 10 independently initialized models it does not reliably reproduce the strong ring-like and asymmetric-connectivity motifs observed in the HRNN. The unit-matched unstructured RNN does not reliably reproduce all layer-specific distributions. These comparisons clarify the contribution of the hierarchical routing and modular input-output structures (Table 4).

**Table 4:** Comparison of neural representations and ring dynamics across the three recurrent architectures. Values are presented as mean *±* SEM. Statistical significance was assessed using FDR-corrected *p*-values. n.s., not significant; * *p <* 0.05; ** *p <* 0.01; *** *p <* 0.001.

| Metric | Statistic | Unstructured HRNN | Multi-module RNN | HRNN |
| --- | --- | --- | --- | --- |
| AHV-tuned units | Layer 1 | 91.8 ± 0.8% | 100.0 ± 0.0% | 99.0 ± 0.2% |
| | Layer 2 | 88.0 ± 2.3% | 84.4 ± 0.8% | 90.2 ± 0.9% |
|  | Difference | L1 > L2 | L1 > L2 | L1 > L2 |
| | $p_{\text{FDR}}$ | 0.152 (n.s.) | $1.05 \times 10^{-7}$ (***) | $2.39 \times 10^{-5}$ (***) |
| Directional MVL | Layer 1 | 0.388 ± 0.077 | 0.599 ± 0.003 | **0.621 ± 0.002** |
| | Layer 2 | 0.480 ± 0.072 | 0.680 ± 0.009 | **0.745 ± 0.004** |
|  | Difference | L2 > L1 | L2 > L1 | L2 > L1 |
| | $p_{\text{FDR}}$ | 0.0283 (*) | $2.39 \times 10^{-5}$ (***) | $1.29 \times 10^{-8}$ (***) |
| Conjunctive grid cells | Layer 3 | 6.7 ± 1.0% | 9.3 ± 0.6% | **16.8 ± 1.1%** |
| | Layer 4 | 4.8 ± 0.4% | 0.5 ± 0.1% | 2.1 ± 0.6% |
|  | Difference | L3 > L4 | L3 > L4 | L3 > L4 |
| | $p_{\text{FDR}}$ | 0.152 (n.s.) | $2.66 \times 10^{-7}$ (***) | $6.61 \times 10^{-6}$ (***) |
| Pure grid cells | Layer 3 | 1.0 ± 0.4% | 8.2 ± 0.5% | 23.0 ± 3.5% |
| | Layer 4 | 6.2 ± 1.4% | 13.5 ± 1.2% | **41.8 ± 2.7%** |
|  | Difference | L4 > L3 | L4 > L3 | L4 > L3 |
| | $p_{\text{FDR}}$ | 0.00282 (**) | 0.00192 (**) | $2.59 \times 10^{-4}$ (***) |
| Layer-1 ring score | | 0.468 ± 0.085 | 0.215 ± 0.011 | **0.850 ± 0.002** |
| Layer-1 asymmetry score | | 0.266 ± 0.052 | 0.139 ± 0.015 | **0.452 ± 0.003** |

### C.5 Model comparison of different readout layer setups and activation functions

As discussed in [28; 23], previous studies have shown that, in single-layer RNNs trained on spatial path integration tasks, the choice of readout configuration (e.g., low-dimensional Cartesian coordinates, Gaussian tuning, or DoS tuning) and activation function (e.g., tanh or ReLU) can substantially influence the emergence of grid cells in hidden units. To investigate how these architectural choices affect grid cell formation in the HRNN, we systematically varied either the readout configuration or the activation function while keeping all other model settings unchanged (Figs. S6–S7).

For the readout configuration, we first examined the effects of varying the number of readout units and their tuning widths (Fig. S5). We then evaluated two alternative readout configurations: (1) a low-dimensional coordinate readout (sin *θ*, cos *θ, x, y*) and (2) Gaussian-tuned direction- and place-cell readout layers (Fig. S6). Finally, to assess the influence of the activation function, we compared the HRNN using tanh activation with the original model (Fig. S7).

In summary, the emergence of direction cells and ring attractor dynamics in Layers 1 and 2 was largely insensitive to variations in readout parameters and readout configurations (coordinate-based or DoS). Similarly, ring attractor dynamics emerged in both layers across all activation function configurations, with the exception of Layer 1 when using tanh activation. In contrast, the emergence of grid cells in Layers 3 and 4 was sensitive to both the number of place-cell readout units and their tuning widths. Moreover, grid cells emerged only when DoS readout and ReLU activation were used, consistent with previous findings [28; 23].

**Figure S5:**
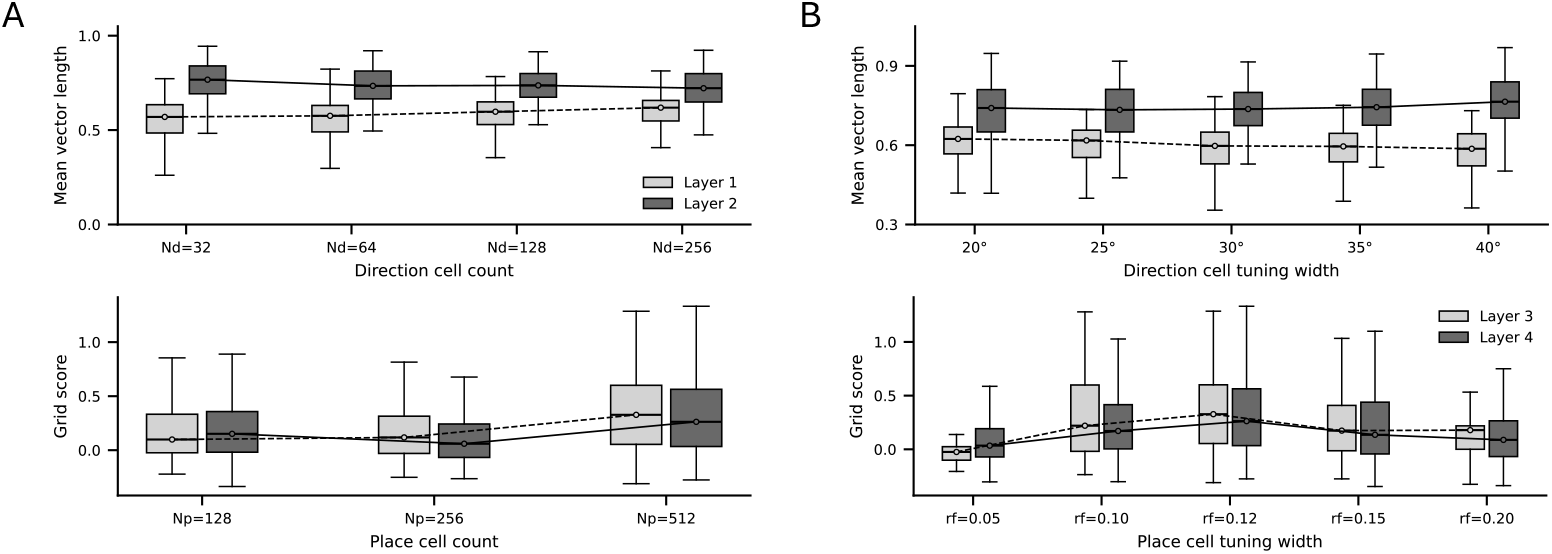
Sensitivity of cell-type properties to readout-neuron tuning parameters. (A) Upper: Mean vector length of directional tuning for different numbers of directional readout cells. Lower: Grid scores for different numbers of place readout cells. (B) Upper: Mean vector length of directional tuning for different tuning widths of directional readout cells. Lower: Grid scores for different tuning widths of place readout cells.

**Figure S6:**
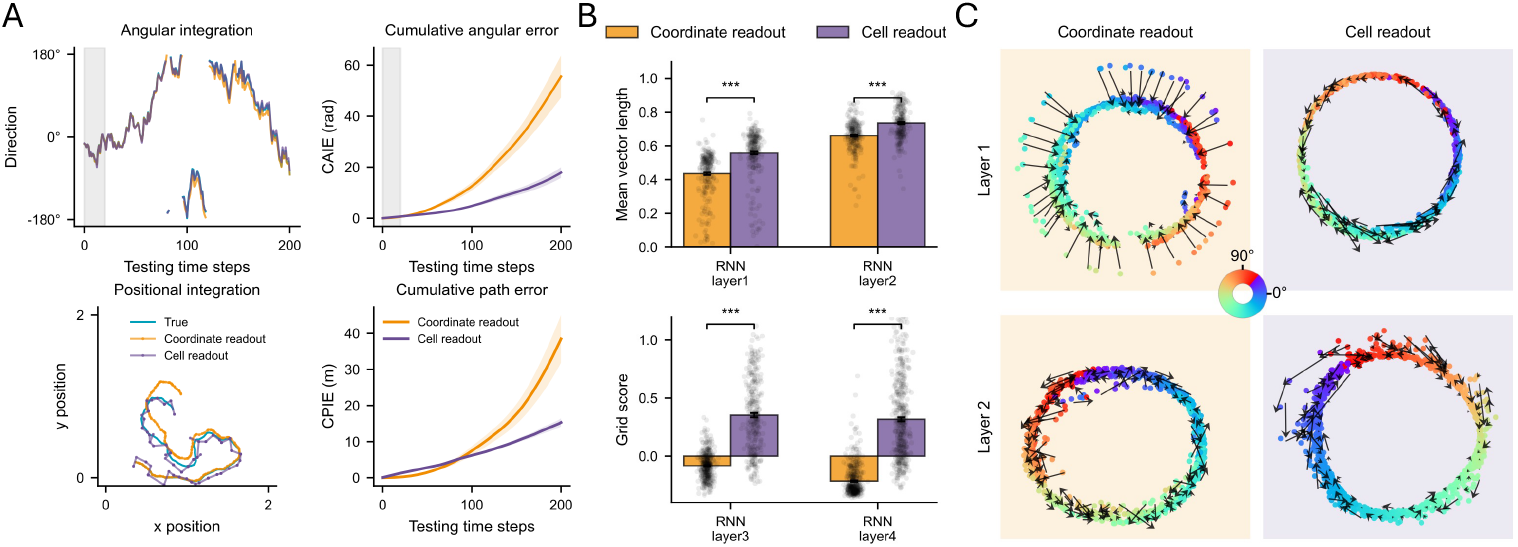
Performance of models with different readout setups. (A) Path-integration performance in models with (1) coordinate readout (sin *θ*, cos *θ, x, y*) and (2) direction- and place-cell readout. (B) Upper: Mean vector length of directional tuning in layers 1 and 2 in the two models. Lower: Grid scores in layers 3 and 4 in the two models. (C) Ring-attractor manifolds in layer 1 (upper) and layer 2 (lower) in the two models.

**Figure S7:**
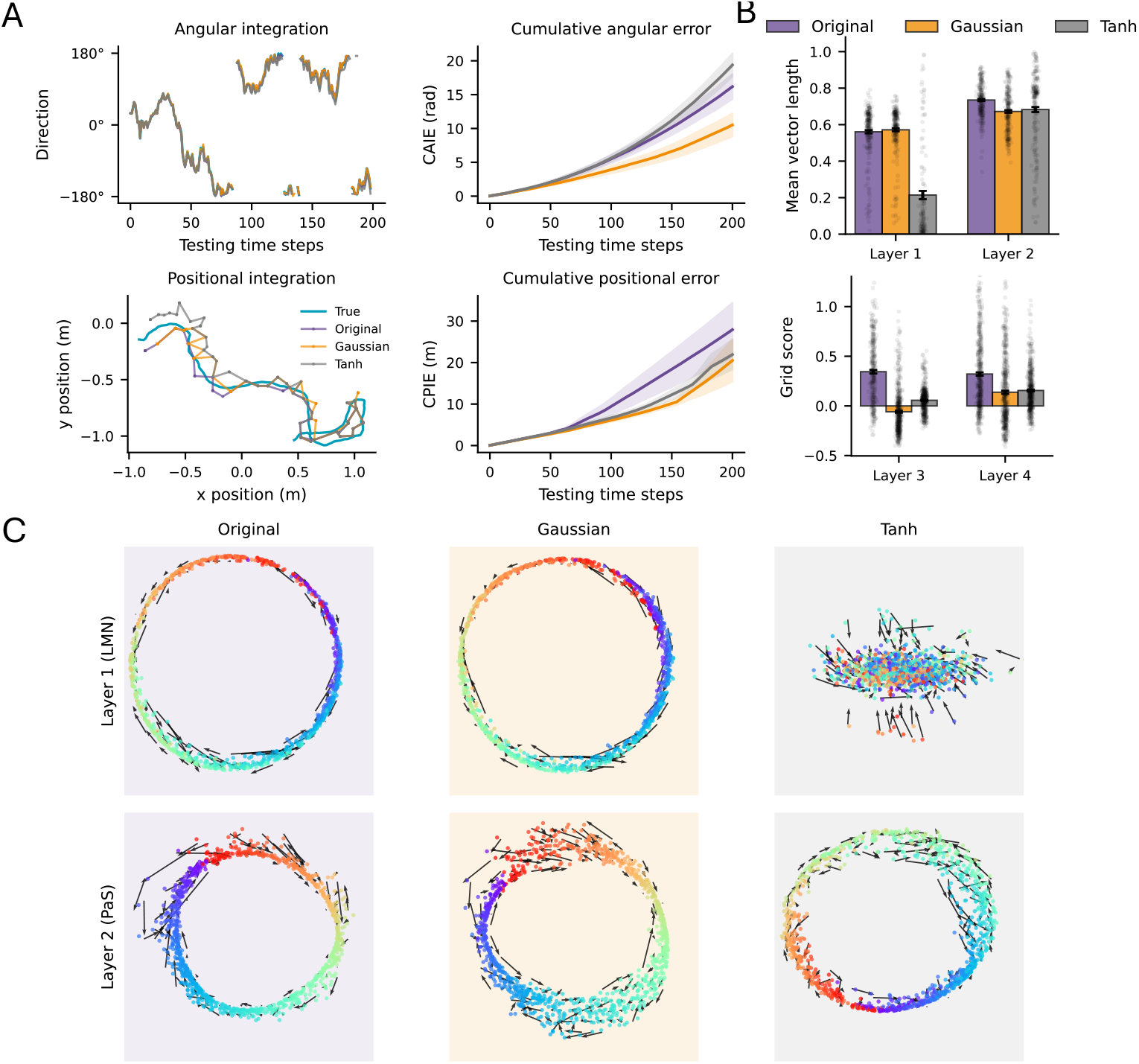
Performance of models with different place-cell readout patterns and activation functions. (A) Path-integration performance in models with (1) DoS readout and ReLU activation, (2) Gaussian readout and ReLU activation, and (3) DoS readout and tanh activation. (B) Upper: Mean vector length of directional tuning in layers 1 and 2 in the three models. Lower: Grid scores in layers 3 and 4 in the two models. (C) Ring-attractor manifolds in layer 1 (upper) and layer 2 (lower) in the three models.

## D Criteria for identifying angular velocity cells

For each unit in layer 1 and 2, we first computed an AV tuning curve by binning signed angular velocity and averaging the unit activity within each bin. Let *r*_*i*_ be the mean activity in AV bin *i*, and let *v*_*i*_ be the corresponding AV bin center. Activity values were rectified as

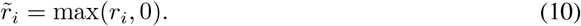

The AV tuning index was defined as a signed modulation depth:

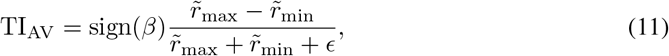

where

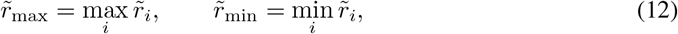

and *ϵ* = 10^*−*8^. The sign was determined by the slope *β* of a linear fit to the AV tuning curve:

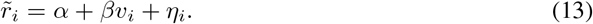

To quantify how well the tuning curve was described by a linear relationship, we computed

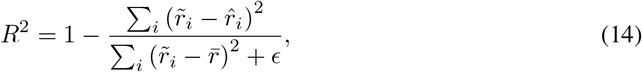

where 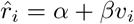 is the fitted response and 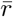 is the mean of 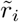 across AV bins. A unit was classified as AV-tuned if

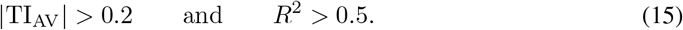

## F Calculating directional mean vector length

Directional tuning was quantified by calculating the mean vector length (MVL). Ground-truth angles were divided into 60 equally spaced bins between *−π* and *π*, and the mean activity of each cell was calculated within each direction bin to obtain a direction-tuning curve. The tuning curves were circularly smoothed using a Gaussian filter with wrap-around boundary conditions.

Only non-silent units were included in the analysis, defined as cells whose peak activity exceeded 10^*−*8^. Each direction-tuning curve was normalized by its total activity across direction bins before computing the MVL. For each cell, MVL was calculated as the magnitude of the activity-weighted circular mean vector:

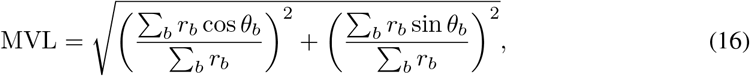

where *r*_*b*_ denotes the normalised mean activity of the cell in direction bin *b* after setting negative values to zero, and *θ*_*b*_ is the centre of that direction bin. Larger MVL values indicate stronger directional modulation. Hidden units with MVL *>* 0.3 were classified as direction-tuned cells, unless otherwise stated.

## F Calculating the grid score

Grid score was calculated following Banino et al. [1]. Only non-silent units with maximum activation exceeding 10^*−*8^ were included in the calculation.

The two-dimensional trajectory arena was first divided into 20 *×* 20 spatial bins. At each time step, the ground-truth position was assigned to the corresponding spatial bin, and activity was averaged across all visits to each bin to obtain a spatial rate map for each unit. For each non-silent cell, the spatial autocorrelogram (SAC) of its spatial rate map was computed. To quantify six-fold rotational symmetry, the SAC was rotated by 30^*°*^, 45^*°*^, 60^*°*^, 90^*°*^, 120^*°*^, 135^*°*^, and 150^*°*^.

Correlations between the original and rotated SACs were calculated within annular masks, excluding the central region of the SAC. For each mask, the grid score compared correlations at grid-aligned rotations (60^*°*^ and 120^*°*^) with those at non-grid-aligned rotations (30^*°*^, 90^*°*^, and 150^*°*^):

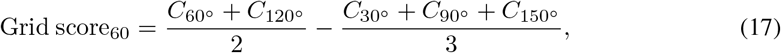

where *C*_*α*_ denotes the correlation between the SAC and the SAC rotated by angle *α* within the annular mask. The final grid score of each unit was defined as the maximum Grid score_60_ across all tested annular masks. Larger values indicate stronger hexagonal grid-like spatial periodicity. Cells with grid score *>* 0.3 were classified as grid cells, unless otherwise stated. Among grid cells, cells with MVL *>* 0.3 were further classified as conjunctive grid cells, whereas those with HD MVL ≤ 0.3 were classified as pure grid cells.

## G Calculating flux in ring attractor dynamics

To visualise the low-dimensional structure in layer 1 and 2, principal component analysis (PCA) was first applied to the population activity in layer 1 and layer 2, separately. The first two principal components were used to visualize the population activity in the low dimensional space.

To further assess whether the observed manifold exhibited attractor-like dynamics, flux analysis was then performed in the PCA-projected state space. This analysis follows the population-manifold approach used by Chaudhuri et al. [6], where local flow fields were examined to determine whether neural states away from the manifold showed biased dynamics back toward the manifold. Here, flux was estimated from temporal transitions in the low-dimensional PCA trajectories of the HRNN hidden states.

To estimate the local flow field, for each time step, the position in PCA subspace was defined as the current hidden state, and the displacement vector was defined as the difference between the next and current PCA states:

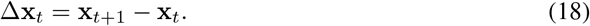

The two-dimensional PCA subspace was divided into 20 *×* 20 bins. Each transition was assigned to a bin according to its starting position **x**_*t*_, and the displacement vectors within each bin were averaged. The resulting mean displacement vector in each occupied bin was defined as the local flux vector:

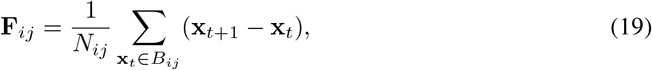

where *B*_*ij*_ denotes the (*i, j*)-th bin in PCA space, and *N*_*ij*_ is the number of transitions assigned to that bin. Only bins containing at least one transition were included. The flux vectors therefore summarize the average direction and magnitude of local hidden-state dynamics in the PCA-projected state space.

## H Calculating offset from layer 3 to layer 4

The spatial offset from layer 3 to layer 4 was quantified using a signed, weight-based centre-of-mass (CoM) analysis. For each cell in layer 3 and layer 4, a spatial CoM was first computed from its two-dimensional spatial rate map.

For the *j*_*th*_ direction cell in layer 3 (MVL > 0.5), its outgoing weights to the *i*_*th*_ cell in layer 4 were extracted: *W*_*ij*_.

The signed weighted population CoM in layer 4, relative to cell *j* in layer 3, was then computed as:

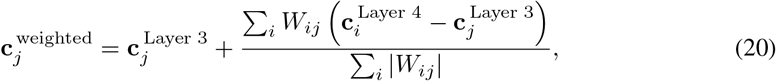

where 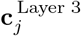 is the spatial CoM of cell *j* in layer 3, 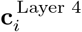 is the spatial CoM of cell *i* in layer 4. Positive and negative feedforward weights shift the weighted population CoM in opposite directions, while normalization by *i ∑*_*i*_|*W*_*ij*_| preserved the signed contribution of each projection.

The spatial offset from layer 3 to layer 4 was defined as the vector from the layer 3 CoM to the signed weighted layer 4 CoM:

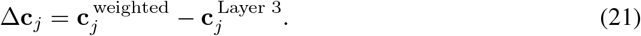

## I Tuning maps of all RNN units

**Figure S8:**
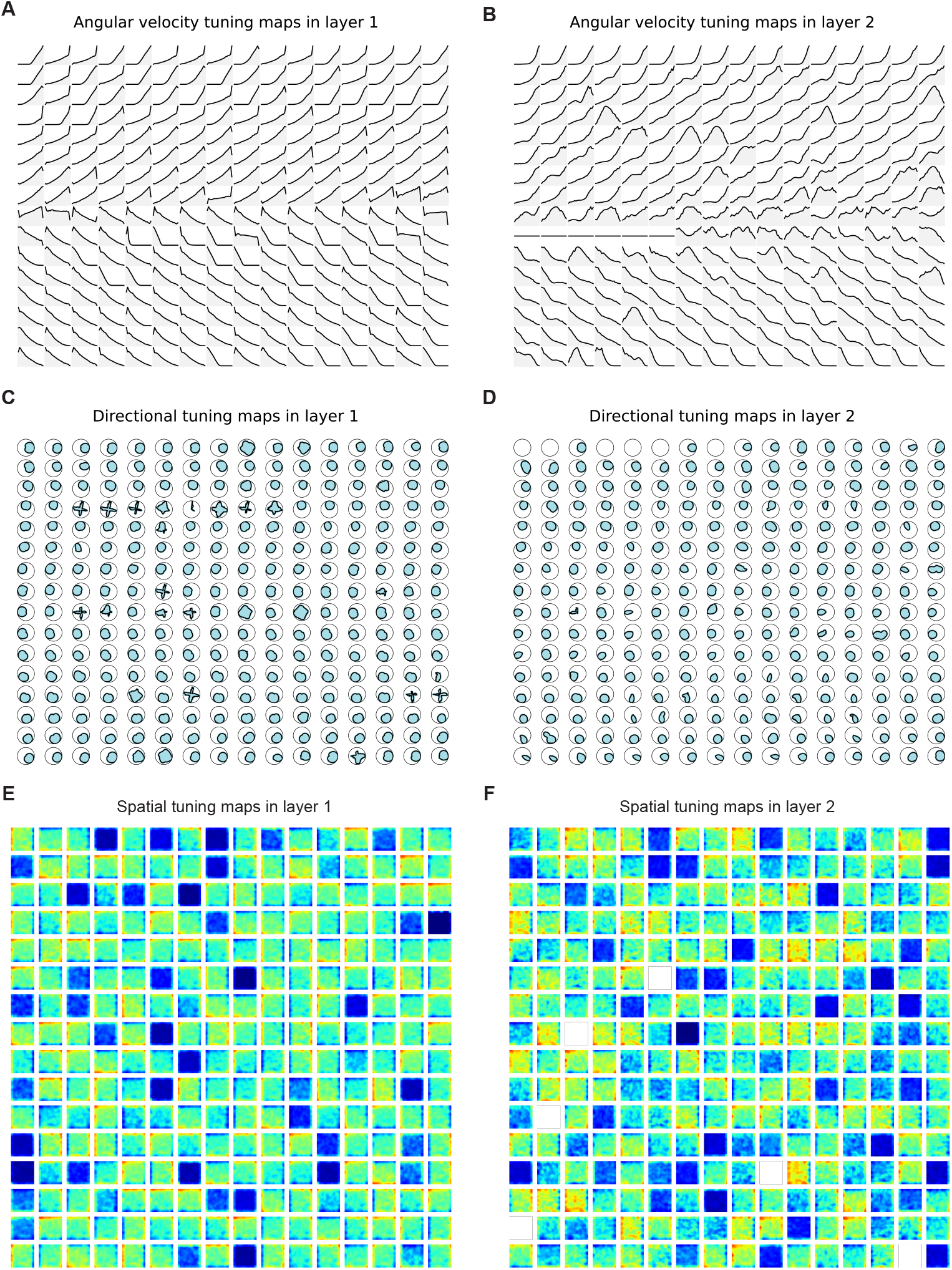
All tuning maps in RNN layers 1 and 2. (A-B) Angular velocity tuning maps in layer 1 and layer 2. (C-D) Directional tuning maps. (E-F) Spatial tuning maps. Note that there is no clear spatial tuning in the first two layers.

**Figure S9:**
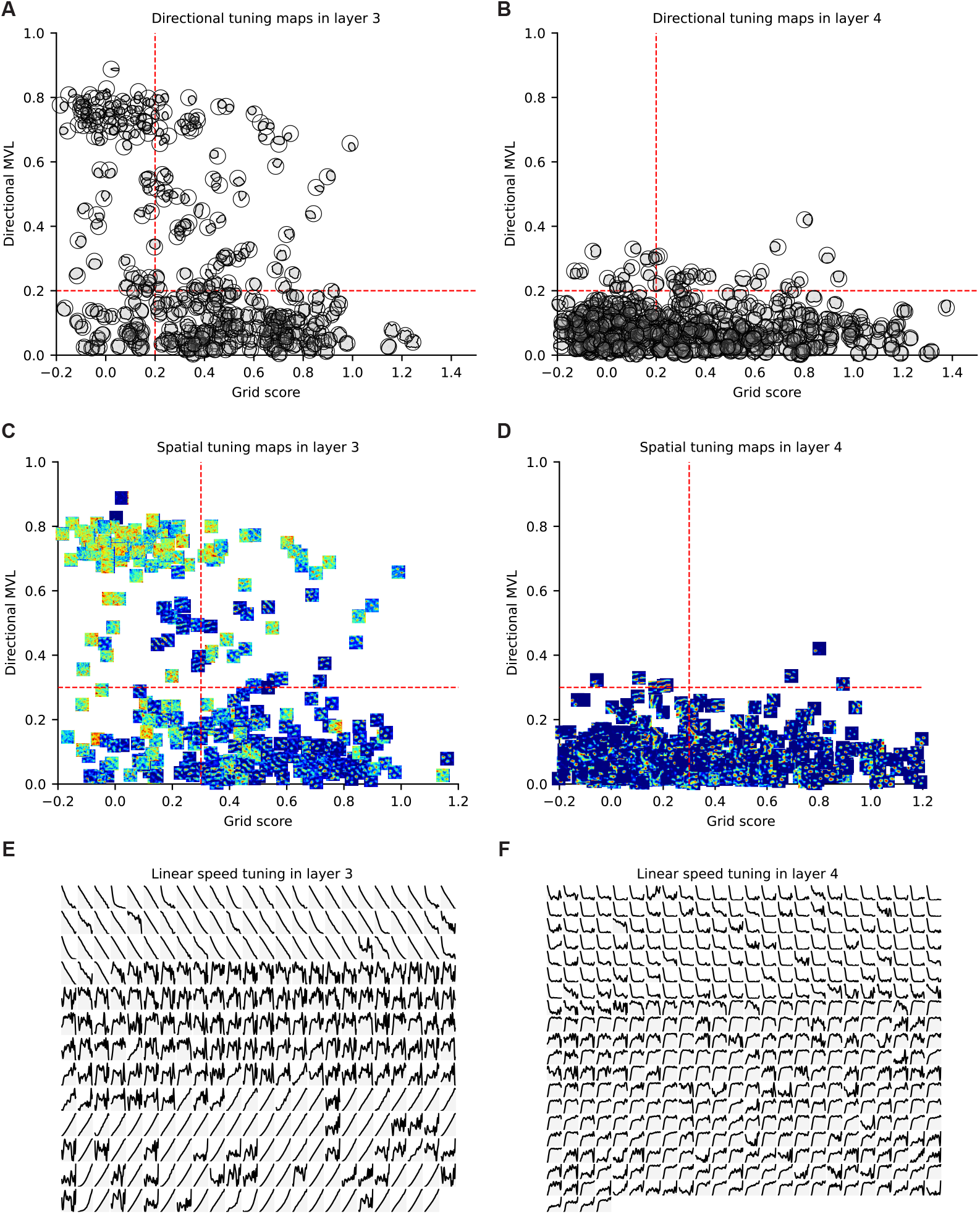
All tuning maps in RNN layer 3 and 4. (A-B) Directional tuning maps in layer 3 and 4, arranged according to each cell’s directional mean vector length (y-axis) and grid score (x-axis). (C-D) Spatial tuning maps in layer 3 and 4, arranged in the same way. (E-F) Linear speed tuning maps in layer 3 and 4, arranged according to the slope of speed tuning from negative to positive. Note that linear speed tuning is more linear in layer 3 compared to layer 4.

